# Impact of Phosphorylation on the Physiological Form of Human alpha-Synuclein in Aqueous Solution

**DOI:** 10.1101/2023.03.10.531864

**Authors:** Emile de Bruyn, Anton Emil Dorn, Giulia Rossetti, Claudio Fernandez, Tiago F. Outeiro, Jörg B. Schulz, Paolo Carloni

## Abstract

Serine 129 can be phosphorylated in pathological inclusions formed by the intrinsically disordered protein human *α*-synuclein (AS), a key player in Parkinson’s disease and other synucleinopathies. Here, molecular simulations provide insight into the structural ensemble of phosphorylated AS. The simulations suggest that phosphorylation does not impact the structural content of the physiological AS conformational ensemble in aqueous solution, as the phosphate group is mostly solvated. The hydrophobic region of AS contains *β*-hairpin structures, which may increase the propensity of the protein to undergo amyloid formation, as seen in the non-physiological (non-acetylated) form of the protein in a recent molecular simulation study. Our findings are consistent with existing experimental data, with the caveat of the observed limitations of the force field for the phosphorylated moiety.

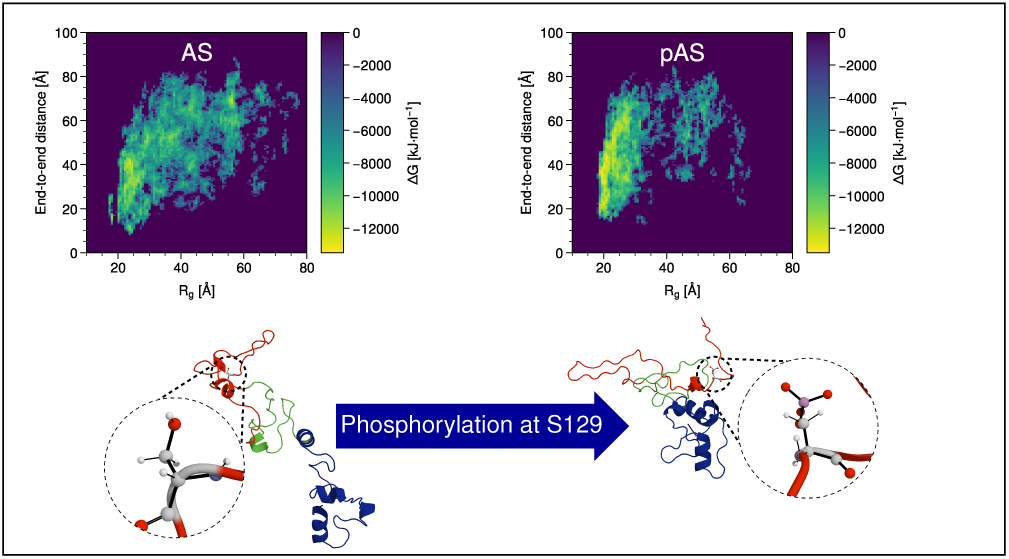

## Introduction

Parkinsons disease (PD) is the second most common neurodegenerative disease after Alzheimer’s disease,^1^ affecting several million people worldwide.^2,3^ The typical pathological hallmark is the accumulation of fibrillar protein inclusions, know as Lewy bodies (LBs) and Lewy neurites (LNs) in the brain.^4,5^ The major component of LBs and LNs are fibrillar forms of the human *α*-synuclein (AS) protein.^3,6^ AS is a 140 amino acid ’disordered’ conformational ensemble both in aqueous solution and in vivo. AS acquires some degree of structure when bound to the membrane or to cellular partners.^7,8^

The primary sequence of AS can be divided in three domains: the positively charged N-terminus (residues 1-60), the overall neutral hydrophobic region (residues 61-95)^1^ and the negatively charged C-terminal domain (residues 96-140, Figure 1). Under physiological conditions, the protein is acetylated on the first residue^2^ In LBs, a significant fraction of AS is phosphorylated on S129.^3 11^ S129 phosphorylation may be regulated by neuronal activity, suggesting that the process may be part of the normal physiology of AS.^12,13^ This post-translational modification (PTM) might play also a pathological role.^14–17^ S129 phosphorylation may be regulated by neuronal activity, suggesting that the process may be part of the normal physiology of AS.^12,13^

**Figure 1:**
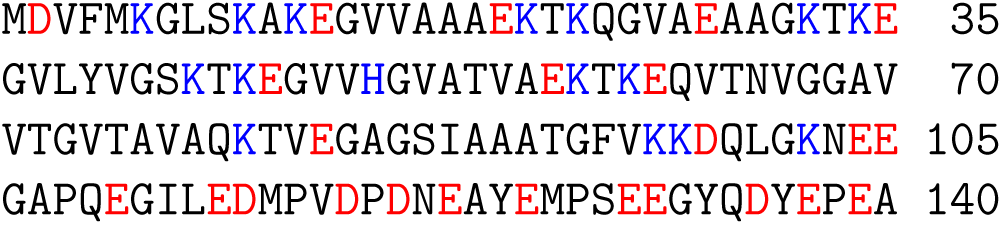
Sequence of amino acid residues in AS; positively charged residues are highlighted in blue and negatively charged ones in red. Three domains can be identified: the positively charged N-terminus (residues 1-60), the overall neutral hydrophobic region (residues 61-95) and the negatively charged C-terminal domain (residues 96-104). In physiological conditions, the protein is acetylated on the first residue, although this post-translational modification does not significantly affect the fibrillization propensity in vitro.^18^ In LBs, a significant fraction of AS is phosphorylated on S129.^11^ A novel phosphorylation site at T64 has also been recently described.^10^

The formation of the 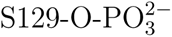 group at the C-terminus of phosphorylated *α* synuclein (pAS) instead of one of the other two domains is intriguing, because it introduces as many as two negative charges at physiological pH (the pKa_1_ and pKa_2_ of phosphoserine are *<* 2 and 5.6^19^).^4^

The impact of phosphorylation on the structural ensemble and aggregation propensity of physiological AS is not known. Thus far, Circular Dichroisim (CD) studies on the non-N-term acetylated form of the protein show that the conformational ensemble does not change significantly upon S129 phosphorylation in solution.^20,21^^5^ On the detailed molecular level, replica exchange simulations based on the CHARMM36m force field^23^ point to an increase of looped secondary structure close to a *β*-hairpin spread throughout the hydrophobic region upon phosphorylation.^24^ However, the structure of the physiological form differs from that of the non-acetylated one (which does not exist in human cells),^25–27^ so firm conclusions on the effect of phosphorylation on endogenous AS cannot be made from these studies.

Here we investigate the impact of phosphorylation on AS’ physiological form by molecular simulation. For this study, one may face several challenges. First, the force field must be adequate to describe IDPs such as AS. The DES-Amber ff99SB,^28^ the Amber a99SB-*disp* ^29^ and CHARMM36m^23^ force fields have been tailored for IDPs^29–31^ and the last two have already been successfully used for the non-acetylated form of the proteins. ^29,31,32^ All of these force fields appear therefore to be well suited to study AS. Second, accurately describing a doubly charged group such as phosphate in pAS is non-trivial. Parameters for phosphorylated serine, threonine, and tyrosine for the Amber^33–35^ and CHARMM^36,37^ force fields have been published. Those for the DES-Amber force field have been recently calibrated on osmotic coefficient calculations.^38^ Unfortunately, simulation studies based on such parametrizations have at times shown artifacts.^39–42^ In addition, none of them describe the hydration properties of the phosphate moiety.^39–47^ This may be very important to fully establish the accuracy of the parametrization. Finally, the conformational space of the protein structural ensemble needs to be efficiently explored. Among the many methodologies used to investigate IDPs successfully,^48–57^ our predictions based on Replica Exchange with Solute Tempering 2 (REST2)^58^ enhanced sampling predictions of wild-type^59,60^ and mutants of AS,^60,61^ turned out to reproduce a variety of biophysical properties of the protein and hence they appear well suitable to study this problem.

Here, we present REST2 simulations of AS and pAS based on the DES-Amber^38^ and a99SB-*disp* force-fields.^29^ We use TIP4P-D for DES-Amber, and the accompanying modified TIP4P-D water model for a99SB-*disp*. To the best of the authors’ knowledge, these simulations are the only ones so far (i) reporting on the physiological form of AS in explicit solvent,^6^ and (ii) describing in detail the hydration properties of the phosphate.

## Methods

### Molecular Simulations

The initial structure of the acetylated protein (AS) which best reproduced the chemical shifts in ref. 86 was selected from the conformational ensemble previously reported in ref. 59. The phosphorylated protein (pAS) was built by adding a phosphate group to S129 using PyMOL.^87^

AS and pAS were inserted in a water-filled dodecahedral simulation box with periodic boundary conditions and minimum distance of 35 Å between the protein and the box edges. Na^+^ and Cl*^−^* ions were added to neutralize the system and achieve a concentration of 150 mmol L^−1^. Table 1 shows the composition of the systems.

**Table 1:** Number of atoms in the simulation box for the three systems simulated here.

|  | Protein | Water | Sodium | Chlorine |
| --- | --- | --- | --- | --- |
| AS | 2,020 | 190,533 | 186 | 176 |
| pAS | 2,023 | 172,359 | 171 | 159 |

The simulations were based on: (i) the DES-Amber force field^28^ and the standard TIP4P-D water model^88^ (Table S1 for a full list of the parameters used); (ii) the a99SB-*disp* force field^29^ and its accompanying modified TIP4P-D water model. ^29^ Long range electrostatics were evaluated using the Particle-Mesh Ewald (PME) method, ^89^ using a cutoff distance of 12 Å in real space. The van der Waals interactions featured the same cutoff.

Constant temperature conditions were achieved by coupling the systems with a Nosé-Hoover thermostat^90^ at 300 K, with a time constant of 0.5 ps. Constant pressure was achieved with a Parrinello-Rahman barostat^91^ at 1 bar, with a time constant of 2 ps (Table S1). The LINCS algorithm was used for all bonds involving hydrogen atoms.^92^ The equations of motions were integrated using the md leap-frog algorithm, with a timestep of 2 fs.

The proteins underwent energy minimization (Table S2), and, subsequently 100 ps of MD in the NVT ensemble (Table S3). Then, they were heated up in 25 ps-long steps of 5 K in the same ensemble up to 300 K using simulated annealing (Tables S4 and S5). The systems were further equilibrated for 1 ns in the NPT ensemble (Table S6). Finally, they underwent 100 ns REST2 simulations^58^ in the NPT ensemble. The proteins were not found to be near their periodic images at distances lower than 12 Å during any of these simulations. The simulations converged after 12 ns (see the Results Section).

Structurally similar conformational clusters were obtained following the method for clustering IDPs described in ref. 93: For both AS and pAS, a total of 8.000 frames from the converged trajectories were clustered (Figures S1 and S2).

The following properties were calculated for conformations after convergence: (i) The radius of gyration *R_g_*, calculated using the MDTraj Python code.^94^ (ii) The hydrodynamic radius, calculated from the radii of gyration using the linear fit of ref. 72. (iii) The NMR chemical shifts of the C*_α_* atoms, calculated using the shiftx2 code. ^95^ (iv) The CD spectra, using the DichroCalc code.^96^ Additionally the average of all conformational clusters, weighted to their occurrence along the trajectory, was calculated. (v) The solvent accessible surface area (SASA) using the MDTraj code. ^94^ (vi) The contact map of protein residues using minimum pairwise distances between residues using the MDTraj code.^94^ (vii) Radial distribution functions (RDFs) from the oxygen atoms of S129 using the SPEADI^97,98^ code developed by the authors.

Hydrogen bonds were defined according to the scheme in ref. 99. These were identified using the MDTraj code. ^94^ (ix) Salt bridges were defined using a distance between two charged atoms in the protein at a distance below 3.25 Å as in ref. 100. The MDTraj code^94^ was also used to identify these. Secondary structure elements were identified using MDTraj^94^ and DSSP.^101^

## Results and Discussion

We performed REST2 simulations^58^ for 100 ns, using 32 replicas, for both AS and pAS in aqueous solution. The calculations were based on the DES-Amber^38^ and the a99SB-*disp* force fields,^29^ both already used for IDPs. We report results at length for calculations using the former, while we provide a summary for the latter here and details in the Supplementary Information.

### Convergence

We calculated two quantities as a function of simulated time to investigate the convergence of the systems (Figures S3 to S6): (i) the running averages of the percentage of secondary structures. In particular, helix structures reached a plateau after 12 ns; (ii) the running averages of the C*_α_* chemical shifts which converge closely to the experimental values within 12 ns. Because of the limitations of the standard usage of RMSD with IDPs such as AS,^102^ the running RMSD of atomic positions was not taken into account. Based on this analysis, we calculated structural properties in the interval 12–100 ns.

### Comparison with experiment

The computational setup results in trajectories that reproduce the experimental data (NMR, CD and hydrodynamic radius)^21,27,86^ available for AS (Figures S5 to S7 and Table S8).

### Effect of phosphorylation

The ten most frequent conformational clusters of AS and pAS represent a total of 47.7 % and 52.9 % of the converged simulation trajectories, respectively (Table S7). They are presented in Figures 2 and 3.

**Figure 2:**
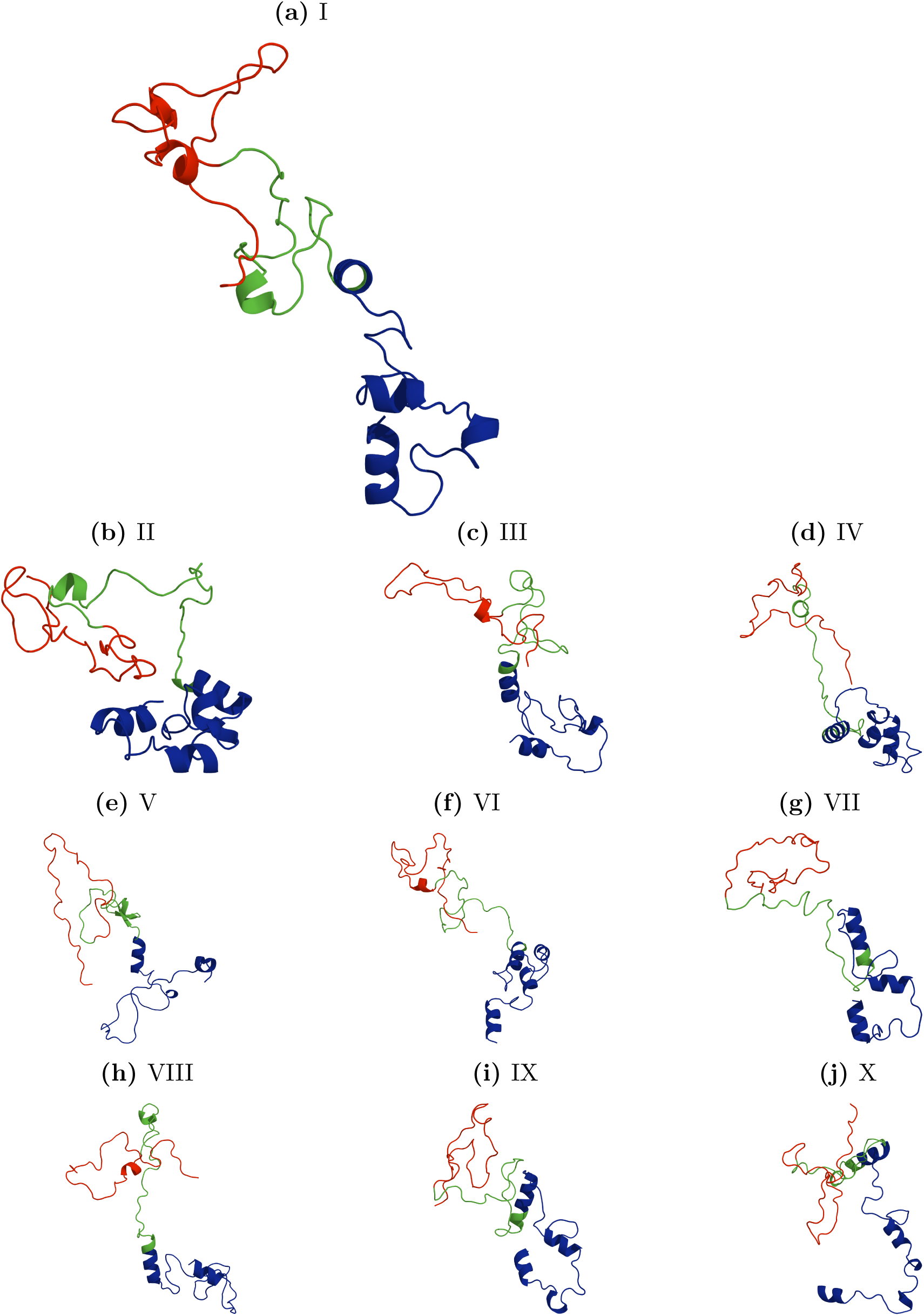
Representative structures (I-X) of AS as determined by clustering. The N-terminal domain is colored blue, the hydrophobic region green and the C-terminal domain red. The percentage of occurrence of the structures decreases from left to right and top to bottom (Table S7).

**Figure 3:**
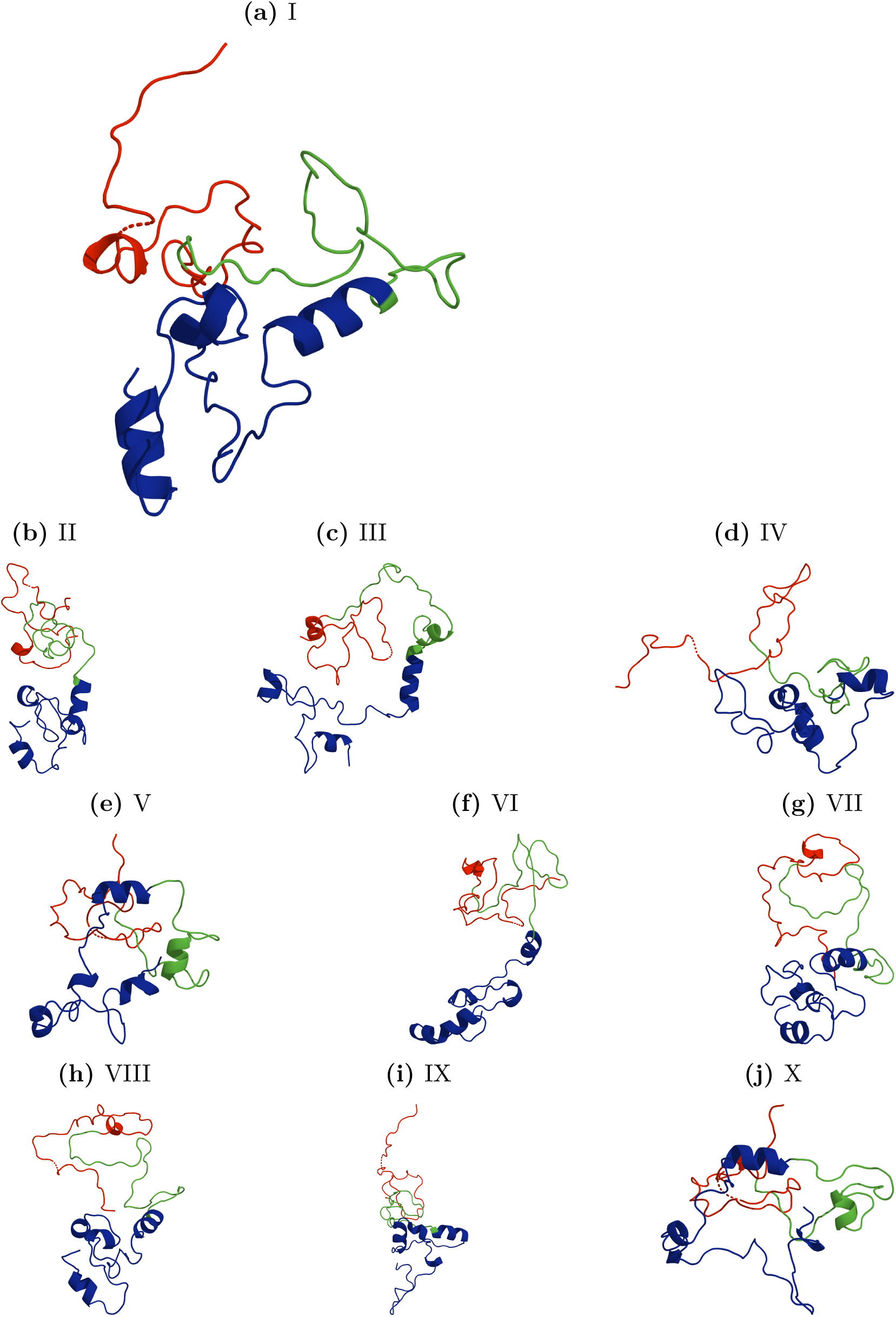
Representative structures (I-X) of pAS as determined by clustering. The acetyl group at the N-terminus and the residue at position S129 are shown as ball and sticks. The percentage of occurrence of the structures decreases from left to right and top to bottom (Table S7).

The mean calculated hydrodynamic radii agree with the experimental values (28.2 and 35.3 Å for AS and pAS,^22^ respectively), within the standard deviation (Table 2). These and the mean radii of gyration (*R_g_*) decrease slightly upon phosphorylation (still within the margin of error, Table 2). The distribution of *R_g_*, however, differs significantly between AS and pAS (Figure 4(a)): the latter displays a higher propensity for either compact or extended conformations than AS. At the qualitative level, this changes the free energy landscape of the protein to form two slightly deeper basins instead of a more shallow landscape (Figures 4(b) and 4(c)). Interestingly, pAS displays a higher likelihood to retain a compact conformation, even when the distance between the termini increases.

**Figure 4:**
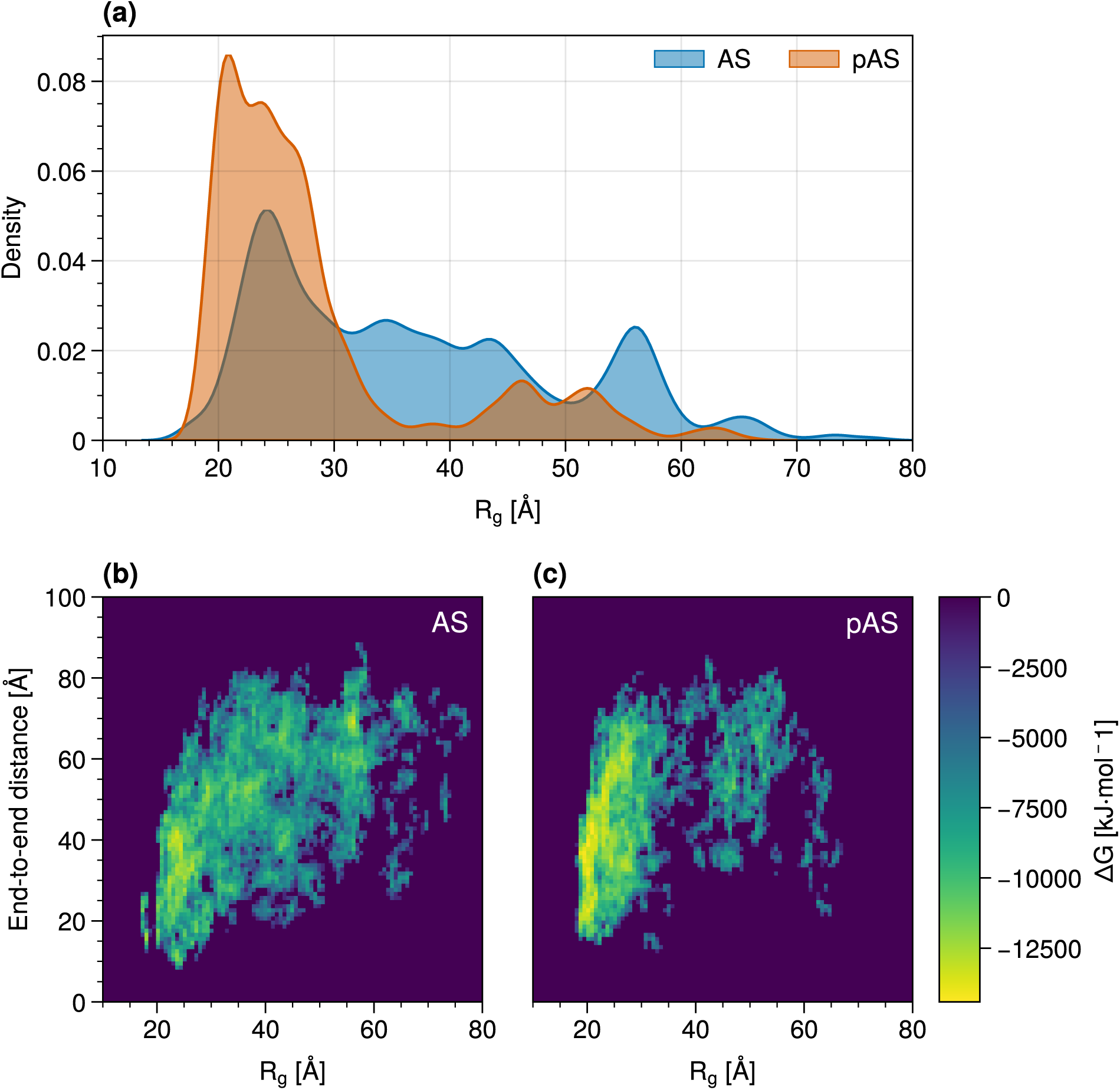
*R_g_* distribution for AS (blue) and pAS (orange) (a) and the corresponding (highly approximate) free energy landscapes plotted as a function of the distance between the protein termini (b-c).

**Table 2:** Calculated properties of AS and pAS. (i) Hydrodynamic *R_H_* and gyration (*R_g_*) radii of the entire proteins and of the hydrophobic regions (HR). (ii) Average number of hydrogen bonds. (iii) Average number of salt bridges. Standard deviations are indicated in parenthesis.

| Protein | $R_H$ [$\text{\AA}$] | $R_g$ [$\text{\AA}$] | $R_g(\text{HR})$ [$\text{\AA}$] | $R_H(\text{HR})$ [$\text{\AA}$] | $N_{SB}$ | $N_{HB}$ |
| --- | --- | --- | --- | --- | --- | --- |
| AS | 35.1 (± 6.1) | 35.6 (± 12.8) | 23.8 (± 16.4) | 28.3 (± 11.3) | 1.54 (± 1.23) | 19.52 (± 4.28) |
| pAS | 31.0 (± 5.8) | 28.0 (± 9.6) | 17.2 (± 9.2) | 23.1 (± 8.1) | 1.86 (± 1.75) | 20.67 (± 4.32) |

The solvent accessibility increases in the hydrophobic region (Table 3). However, because of the large fluctuations of the C-terminus – N-terminus distances (Figure S8), this analysis is not conclusive.

**Table 3:** Average Solvent Accessible Surface Area (SASA) in AS and pAS in the hydrophobic region (HR) and at the N- and C-termini.

| Protein | SASA<sub>N</sub> [$\text{\AA}^2$] | SASA<sub>HR</sub> [$\text{\AA}^2$] | SASA<sub>C</sub> [$\text{\AA}^2$] |
| --- | --- | --- | --- |
| AS | 96 ($\pm 41$) | 87 ($\pm 31$) | 111 ($\pm 32$) |
| pAS | 90 ($\pm 42$) | 91 ($\pm 32$) | 113 ($\pm 39$) |

The intramolecular interactions between the hydrophobic region and the other regions slightly increase upon phosphorylation, as shown by the contact maps for each monomer and their differences (Figure 5). Accordingly, the number of intramolecular hydrogen bonds increases (6(a-b)). However, the results should be taken with caution, as the number of such interactions shows a wide normal distribution with a large standard deviation (Table 2).

**Figure 5:**
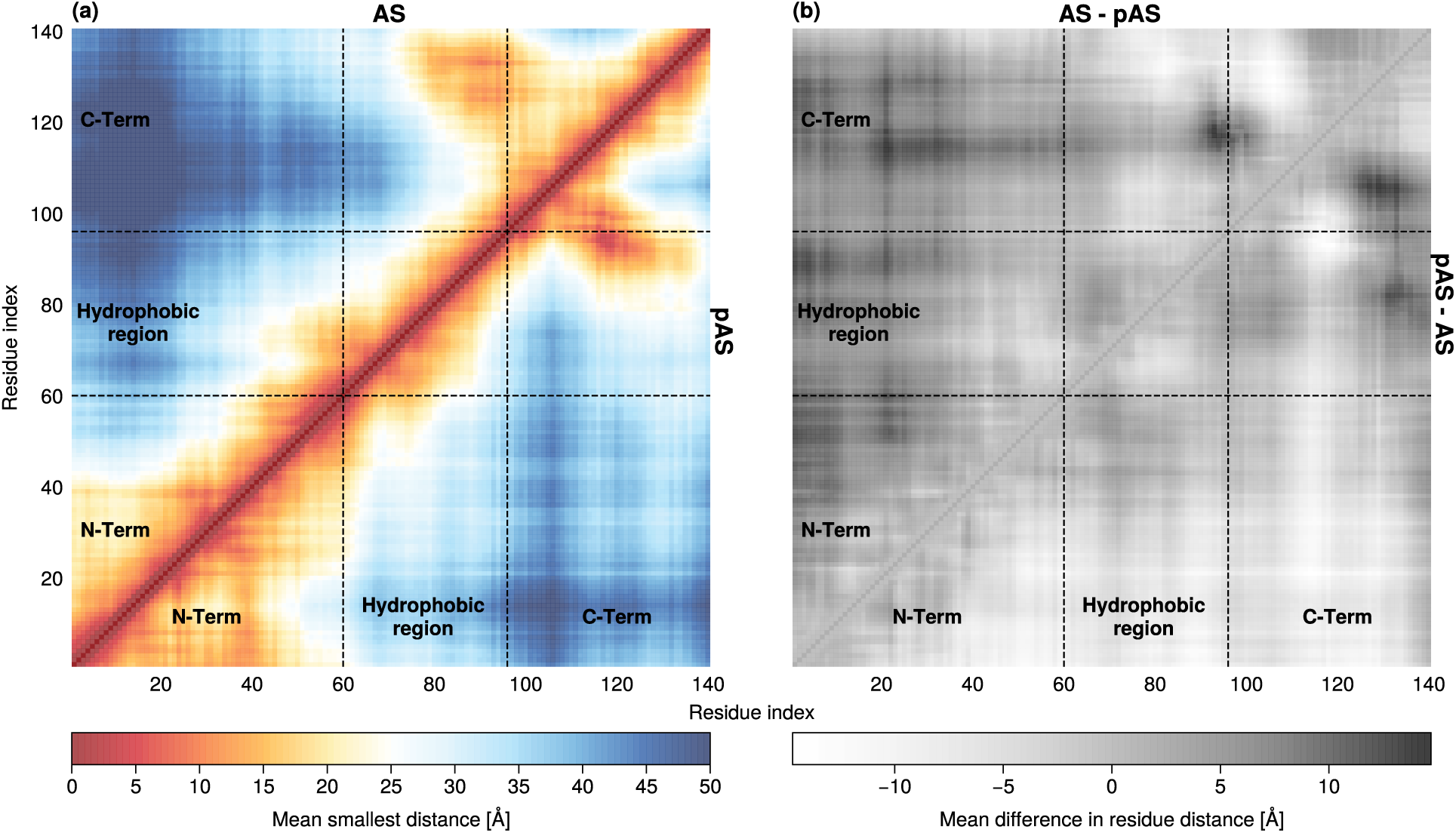
(a) Contact maps of AS (triangle above) and pAS (triangle below). (b) Mean residue distance differences between AS (triangle above) and pAS (triangle below). Brighter corresponds to comparatively closer distances.

These H-bonds are almost exclusively formed within every single domain (Figures 6(a-b)). The number of salt bridges increases upon phosphorylation. The most persistent salt bridges common to both forms involve the N-term and hydrophobic regions (K23–E20 and K58–E61, respectively, see Figures 6(a-b)). Few salt bridges are formed between the C-terminus and one of the two other domains. Of course, because the intramolecular contacts of IDPs are labile, the scenario could be different over longer time scales or at different starting points.

**Figure 6:**
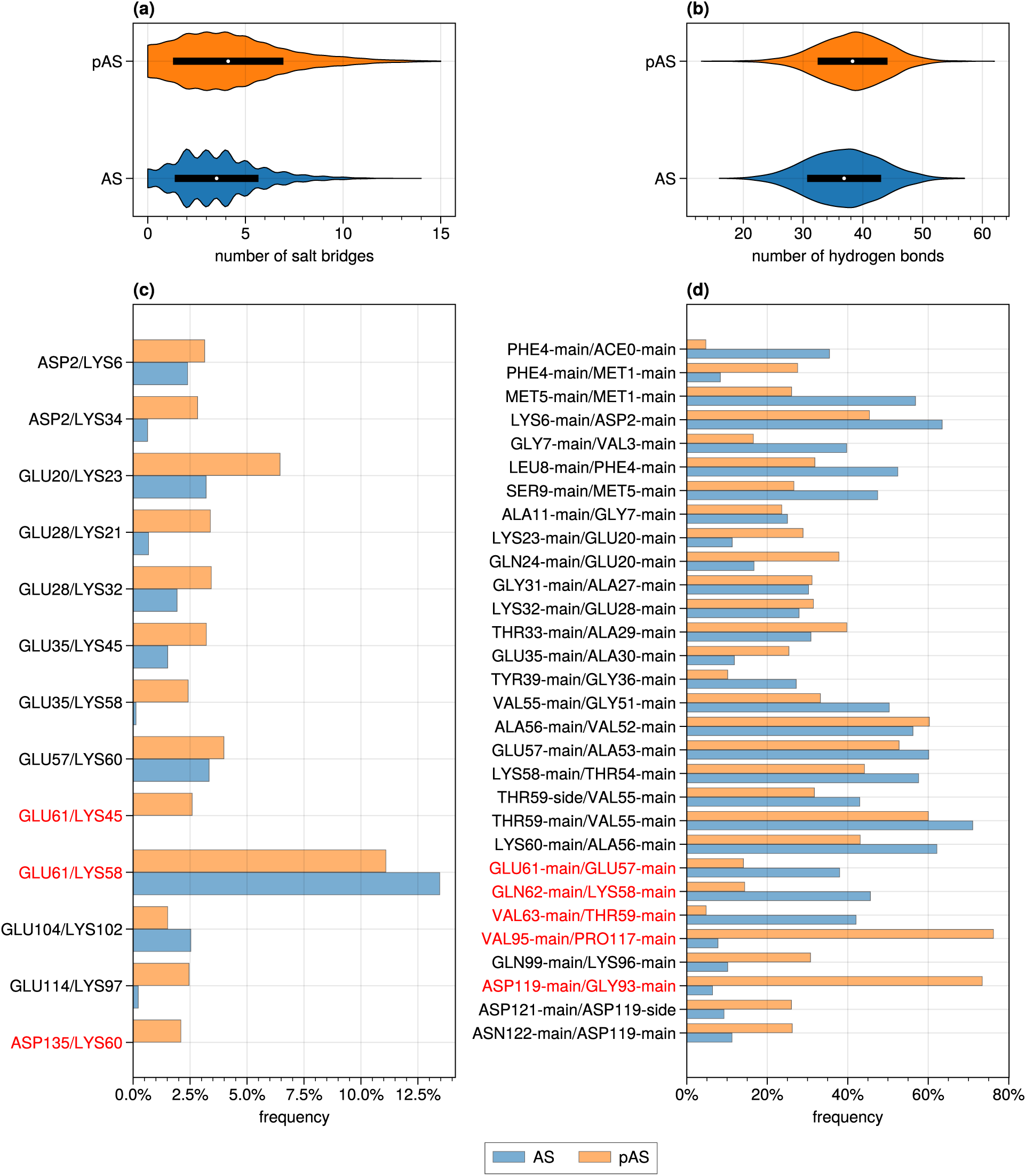
Distribution of the total number of salt bridges (a) and hydrogen bonds (b) in AS (blue) and pAS (orange). Frequency with which intradomain (black labels) and interdomain (red labels) salt bridges (a) and hydrogen bonds (b) are found in AS and pAS. Shown are salt bridges and hydrogen bonds that occur during at least 2 % and 25 % of the converged trajectory, respectively, in either the AS or pAS simulation.

We conclude that the protein’s conformational ensemble becomes *slightly*more compact upon phosphorylation. This is associated with a small increase of intramolecular interactions. This is expected, as the increase of negative charge at the C-terminus can increase its interaction with the rest of the protein, and particularly with the positively charged N-terminus: the phosphate group interacts to a small extent with sodium ions (Figure S17) and, even less, with the protein residues (Figure S16). In fact, this group is fully solvent exposed, as shown by a plot of the phosphate oxygen-water oxygen RDFs (Figure 7). This may be caused, at least in part, by the presence of four negatively charged phosphate oxygen atoms.^7^ Within these caveats, we suggest that the electrostatic field of the phosphate is strongly reduced by its solvation and does not impact the C-terminus-N-terminus interactions. This lack of interaction may also in part be explained by AS containing no arginine residues that are known to interact strongly with phosphate groups,^103^ and indeed none of the lysine residues present in the N-terminal domain formed salt bridges with S129 during the simulation (Figure 6).

**Figure 7:**
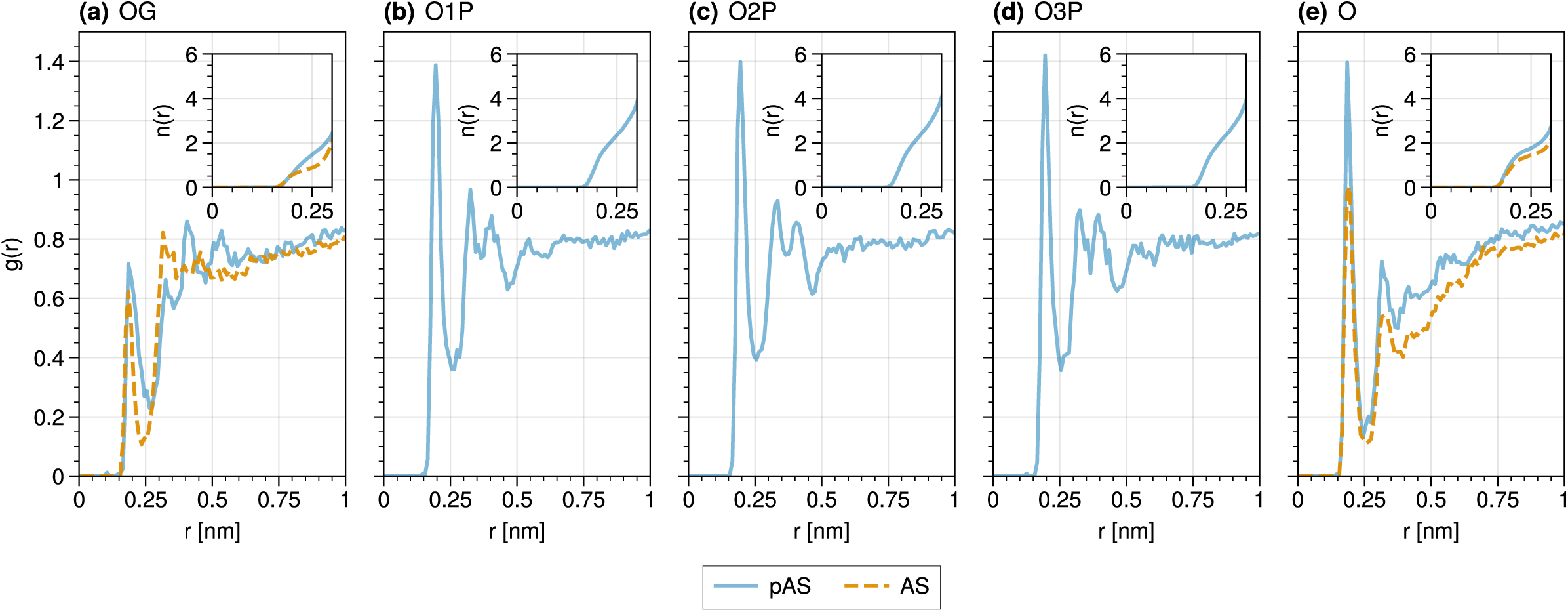
RDFs (*g*(*r*)) of the oxygen atoms of residue 129 and water hydrogens (a-e). Compare the RDF for pAS (blue) and AS (orange) for the oxygen atom in the side chain (a) and the carbonyl oxygen (e). (b-d) illustrate the RDF for the terminal phosphoryl oxygens. Insets show the integral of *g*(*r*) up to 0.3 nm.

In AS, S129 interacts much less with the solvent than phosphate in pAS: The O*γ* atom has only 2 water hydrogen (Figure 7) and no sodium ions in its surrounding. The serine oxydril group instead forms a variety of *intramolecular* H-bonds (with K80, K96, K97, K102, E126, E130 and E131 (Figure S16). S129 backbone units are observed to interact both with the solvent and nearby protein hydrogens in both AS and pAS. Thus, S129 forms many more intermolecular contacts than the phosphorylated residue in pAS.

Using the approach of ref. 24, we find a significant content of *β*-hairpin-like structure in the hydrophobic region upon phosphorylation (Figure S20).^8^ As discussed in ref. 24, these types of structures may be associated with amyloid-forming conformations and hence this finding does suggest that phosphorylation increases fibril formation starting from conformations similar to those found in the fibrils.^104,105^ This would be consistent with the fact that almost all the proteins in the fibrils are phosphorylated *in vivo*.^11^

These simulation results show overall only a small change in the structural ensemble of aqueous physiological AS upon phosphorylation at S129. Our previously published simulations of familial-PD linked mutations of AS^98^ display far greater changes in the structural ensemble relative to the wild-type, compared to phosphorylation in the current study. The simulations here are in line with experimental studies showing that phosphorylation and de-phosphorylation of AS are likely normal physiological processes fine-tuning binding to lipids,^106,107^ not a clear marker of pathology.^12,13^ This, however, does not preclude the prevailing view that phosphorylation is also implicated in fibril formation.^12^

The simulations of AS and pAS with the Amber a99SB-*disp* force field^29^ show very similar results as those presented here, except for the phosphate hydration properties, which turn out to be less accurate than those of the DES-Amber force field (See Supplemementary Information, Sections 4, 5).

The simulations of the protein with monoprotonated phosphate (based on the a99SB-*disp* force field) turn out to be rather similar to those of pAS (see Supplementary Information, Section 6). Thus, we conclude that if such species exist in equilibrium with pAS, they contribute to the protein structural ensemble similarly to pAS.

## Conclusions

Because of the presence *in vivo* of phosphorylated AS, the detailed understanding of the impact of phosphorylation on this protein is important for informing therapeutic strategies aimed at targeting AS in synucleinopathies. Here, we investigated the effects of phosphorylation on the structural ensemble of AS in solution by REST2 simulations. Our simulations of AS are consistent with a plethora of experimental data, as in our previous simulations.^59^ Most importantly, our study of the as yet unexplored physiological form of pAS concludes that it also does not undergo large conformational changes relative to the physiological form of AS. The small changes are consistent with CD studies of phosphorylation on the non-physiological form of AS.^20,21^ The protein simply becomes slightly more compact than the unmodified protein. As seen for the non-physiological form of the protein 24, phosphorylation induces a *β*-hairpin-like, amyloid-forming conformations. The increased propensity towards fibril formation is likely consistent with the fact that about 90 % AS in the LBs is phosphorylated.^11^

## Supporting Information Available

The data that support the findings of this study are available from the corresponding author, GR, upon reasonable request.

Detailed experimental setup, analysis and comparison with results obtained with other force fields are contained in the Supplementary Information (PDF).

Structures of the 30 most frequently found cluster midpoints are included for both AS and pAS (PDB).

## Supporting information

Structural cluster midpoints

## Acknowledgement

TFO is supported by DFG (SFB1286-B8) and under Germanys Excellence Strategy - EXC 2067/1-390729940.

This work was partially performed as part of the Helmholtz School for Data Science in Life, Earth and Energy (HDS-LEE) and received funding from the Helmholtz Association of German Research Centers.

Open Access publication funded by the Deutsche Forschungsgemeinschaft (DFG, German Research Foundation) 491111487.

The authors gratefully acknowledge the Gauss Centre for Supercomputing e.V. (www.gauss-centre.eu) for funding this project by providing computing time through the John von Neumann Institute for Computing (NIC) on the GCS Supercomputer JUWELS at Jülich Super-computing Centre (JSC).^108–111^

The authors gratefully acknowledge insightful discussions with Stefano Piana-Agostinetti regarding the choice of force field parameters, in particular with respect to the hydration of the phosphate group.

## Footnotes

1 We choose to use the more accurate term “hydrophobic region” instead of the historical but inaccurate term “non-amyloid component (NAC)”. ^9^

2 N-terminal acetylation does not significantly change the fibrillization propensity in vitro. ^10^

3 A novel phosphorylation site of *α*-synuclein at T64 has also been recently described. ^10^

4 It might be possible that the phosphate is partially monoprotonated. The effect of protonation is discussed in the Supplementary Information.

5 This contrasts with findings by CD studies for phosphorylation on protein variants. ^20–22^ These point to significant changes in the structural ensemble upon phosphorylation.

6 Many molecular simulation studies, besides those in refs. 29,32, focus on the non-acetylated protein. ^31,62–64,64–85^ Calculations of the protein in implicit solvent are not reported here.

7 The phosphate oxygen charges in the CHARMM ^36,37^ and Amber ^34^ parametrizations, also used to study phopshate PTMs in IDPs, are not too dissimilar to those of used here (−0.565to−0.639 e). In the case of Amber-based simulations of pAS, reported in the Supplementary Information, we also see an overestimation of interacting water molecules.

8 This content is however smaller than that observed for the non-physiological form, see details in the Supplementary Information

## Notes

### Competing Interest Statement

The authors have declared no competing interest.

### Summary of Updates

Updated to reflect the latest developments in the field and studies on alpha-Synuclein. A section discussing the content of beta-hairpin-like structures in the hydrophobic region was added to the discussion and SI.

