## Supplementary Information for "Impact of Phosphorylation on the Physiological Form of Human alpha-Synuclein in Aqueous Solution"

Emile de Bruyn,[†,‡,¶] Anton Emil Dorn,[†,‡,§] Giulia Rossetti,[*,‡,‖,⊥] Claudio Fernandez,[#,@] Tiago F. Outeiro,[△,∇,††] Jörg B. Schulz,[¶,⊥,‡‡] and Paolo Carloni[¶,‖]

† *Equally contributed to this work*
‡ *Jülich Supercomputing Centre (JSC), Forschungszentrum Jülich GmbH, 52425 Jülich, Germany*
¶ *Department of Physics, RWTH Aachen University, 52062 Aachen, Germany*
§ *Faculty of Biology, University of Duisburg-Essen, 45141 Essen, Germany*
‖ *Computational Biomedicine (IAS-5/INM-9), Forschungszentrum Jülich GmbH, 52425 Jülich, Germany*
⊥ *Department of Neurology, RWTH Aachen University, 52074 Aachen, Germany*
### *Max Planck Laboratory for Structural Biology, Chemistry and Molecular Biophysics of Rosario (MPLbioR, UNR-MPINAT). Partner of the Max Planck Institute for Multidisciplinary Sciences (MPINAT, MPG). Centro de Estudios Interdisciplinarios, Universidad Nacional de Rosario, S2002LRK Rosario, Argentina*
@ *Department of NMR-based Structural Biology, Max Planck Institute for Multidisciplinary Sciences, 37077 Göttingen, Germany*
△ *Department of Experimental Neurodegeneration, Center for Biostructural Imaging of Neurodegeneration, University Medical Center Göttingen, 37075 Göttingen, Germany*
∇ *Max Planck Institute for Multidisciplinary Sciences, 37075 Göttingen, Germany*
†† *Translational and Clinical Research Institute, Newcastle University, Newcastle upon Tyne NE1 7RU, United Kingdom*
‡‡ *JARA Brain Institute Molecular Neuroscience and Neuroimaging (INM-11), Research Centre Jülich and RWTH Aachen University, 52074 Aachen, Germany*

#### 1 Simulation parameters

Tables S1 to S6 show the parameters used to initiate the different runs in GROMACS. Tables S2 to S6 describe both the preparation for the simulations using REST2 and the unbiased MD simulations. The parameters contained in table S1 were used for each replica during the REST2 simulation.

Table S1: Parameters used in GROMACS for the production run using REST2 and unbiased MD. Unbiased production runs used the same parameters.

| Parameter | Value |
|---|---|
| integrator | md |
| nsteps | 50,000,000 |
| dt [ps] | 0.002 |
| nstxout | 5000 |
| nstvout | 5000 |
| nstenergy | 1000 |
| nstlog | 500 |
| nstxout-compressed | 10 |
| nstcalcenergy | 100 |
| compressed-x-grps | Protein, Ion |
| energygrps | Protein, Water, Ion |
| continuation | yes |
| constraint_algorithm | lincs |
| constraints | h-bonds |
| lincs_iter | 1 |
| lincs_order | 4 |
| cutoff-scheme | Verlet |
| nstlist | 10 |
| coulombtype | PME |
| pme_order | 4 |
| rlist [nm] | 1.2 |
| rcoulomb [nm] | 1.2 |
| rvdw [nm] | 1.2 |
| fourierspacing [nm] | 0.12 |
| tcoupl | Nose-Hoover |
| tc-grps | Protein, Water, Ion |
| tau_t [ps] | 0.5 |
| ref_t [K] | 300 |
| pcoupl | Parrinello-Rahman |
| pcoupltype | isotropic |
| tau_p [ps] | 2.0 |
| ref_p [bar] | 1.0 |
| compressibility [$bar^{-1}$] | 4.5e-5 |
| DispCorr | EnerPres |
| pbc | xyz |
| gen_vel | no |
| nstcomm | 1 |
| comm-grps | Protein, Water, Ion |

Table S2: Parameters used in GROMACS for the energy minimization.

| Parameter | Value |
|---|---|
| integrator | steep |
| nsteps | 100000 |
| emtol [kJ mol$^{-1}$ nm$^{-1}$] | 10 |
| emstep [nm] | 0.01 |
| nstlog | 1000 |
| nstenergy | 100 |
| energygrps | System |
| cutoff-scheme | Verlet |
| nstlist | 1 |
| rlist | 1 |
| pbc | xyz |
| pme_order | 4 |
| vdwtype | Cut-off |
| vdw-modifier | Potential-shift-Verlet |
| rcoulomb [nm] | 1.2 |
| rvdw [nm] | 1.2 |
| epsilon_r [nm] | 1 |
| fourierspacing [nm] | 0.12 |

Table S3: Parameters used in GROMACS for the NVT equilibration.

| Parameter | Value |
|---|---|
| integrator | md |
| nsteps | 50000 |
| dt [ps] | 0.002 |
| nstxout | 500 |
| nstvout | 500 |
| nstenergy | 500 |
| nstlog | 500 |
| continuation | no |
| constraint_algorithm | lincs |
| constraints | h-bonds |
| lincs_iter | 1 |
| lincs_order | 4 |
| cutoff-scheme | Verlet |
| nstlist | 10 |
| rcoulomb [nm] | 1.2 |
| rvdw [nm] | 1.2 |
| DispCorr | EnerPres |

Continued on next page

Continued from previous page

| Parameter | Value |
|---|---|
| coulombtype | PME |
| pme_order | 4 |
| fourierspacing [nm] | 0.12 |
| tcoupl | Nose-Hoover |
| tc-grps | Protein, Water, Ion |
| tau_t [ps] | 0.5 |
| ref_t [K] | 300 |
| pcoupl | no |
| pbc | xyz |
| gen_vel | yes |
| gen_temp [K] | 300 |
| gen_seed | -1 |
| Define | -DPOSRES |

Table S4: Parameters used in GROMACS for the simulated annealing.

| Parameter | Value |
|---|---|
| integrator | md |
| nsteps | 550000 |
| dt [ps] | 0.002 |
| nstxout | 2000 |
| nstvout | 2000 |
| nstenergy | 1000 |
| nstlog | 1000 |
| nstxout-compressed | 5000 |
| compressed-x-grps | System |
| energygrps | System |
| continuation | no |
| constraint_algorithm | lincs |
| constraints | none |
| lincs_iter | 1 |
| lincs_order | 4 |
| lincs-warnangle | 30 |
| morse | no |
| cutoff-scheme | Verlet |
| nstlist | 1 |
| coulombtype | PME |
| pme_order | 4 |
| vdwtype | Cut-off |
| vdw-modifier | Potential-shift-Verlet |

Continued on next page

Continued from previous page

| Parameter | Value |
|---|---|
| rcoulomb [nm] | 1.2 |
| rvdw [nm] | 1.2 |
| epsilon_r [nm] | 1 |
| fourierspacing [nm] | 0.12 |
| tcoupl | Nose-Hoover |
| tc-grps | System |
| tau_t [ps] | 0.1 |
| ref_t [K] | 300 |
| pcoupl | Parrinello-Rahman |
| pcoupltype | isotropic |
| tau_p [ps] | 0.5 |
| ref_p [bar] | 1.0 |
| compressibility [$bar^{-1}$] | 4.5e-5 |
| nstcomm | 1 |
| comm-mode | linear |
| comm-grps | Protein, Water, Ion |
| pbc | xyz |
| DispCorr | EnerPres |
| gen_vel | yes |
| gen-temp | 2 |
| gen-seed | 173529 |
| annealing | single |
| annealing_npoints | 40 |
| Define | -DPOSRES |

Table S5: Temperature and time steps in the simulated annealing run.

| annealing_time | annealing_temp |
|---|---|
| 0 | 2 |
| 50 | 5 |
| 75 | 10 |
| 100 | 15 |
| 125 | 20 |
| 150 | 25 |
| 175 | 30 |
| 200 | 35 |
| 225 | 40 |
| 250 | 45 |
| 275 | 50 |
| 300 | 55 |

Continued on next page

Continued from previous page

| annealing_time | annealing_temp |
|---|---|
| 325 | 60 |
| 350 | 65 |
| 375 | 70 |
| 400 | 75 |
| 425 | 80 |
| 450 | 85 |
| 475 | 90 |
| 500 | 100 |
| 525 | 110 |
| 550 | 120 |
| 575 | 130 |
| 600 | 140 |
| 625 | 150 |
| 650 | 160 |
| 675 | 170 |
| 700 | 180 |
| 725 | 190 |
| 750 | 200 |
| 775 | 210 |
| 800 | 220 |
| 825 | 230 |
| 850 | 240 |
| 875 | 250 |
| 900 | 260 |
| 925 | 270 |
| 950 | 280 |
| 975 | 290 |
| 1000 | 300 |

Table S6: Parameters used in GROMACS for the NPT equilibration.

| Parameter | Value |
|---|---|
| integrator | md |
| nsteps | 500000 |
| dt [ps] | 0.002 |
| nstxout | 500 |
| nstvout | 500 |
| nstenergy | 500 |
| nstlog | 500 |
| continuation | yes |

Continued on next page

Continued from previous page

| Parameter | Value |
|---|---|
| constraint_algorithm | lincs |
| constraints | h-bonds |
| lincs_iter | 1 |
| lincs_order | 4 |
| cutoff-scheme | Verlet |
| nstlist | 10 |
| rcoulomb [nm] | 1.2 |
| rvdw [nm] | 1.2 |
| DispCorr | EnerPres |
| coulombtype | PME |
| pme_order | 4 |
| fourierspacing [nm] | 0.12 |
| tcoupl | Nose-Hoover |
| tc-grps | Protein, Water, Ion |
| tau_t [ps] | 0.5 |
| ref_t [K] | 300 |
| pcoupl | Parrinello-Rahman |
| pcoupltype | semiisotropic |
| tau_p [ps] | 2.0 |
| ref_p [bar] | 1.0 |
| compressibility [bar$^{-1}$] | 4.5e-5 |
| refcoord_scaling | com |
| pbc | xyz |
| gen_vel | no |
| Define | -DPOSRES |

### 2 Obtaining Structurally Similar Clusters

High-dimensional pair-wise root mean-square displacement (RMSD) values between individual trajectory frames were projected onto a lower-dimensional plane using t-distributed Stochastic Neighbor Embedding (t-SNE), following the method described by Appadurai et al.[1] This projection was then K-means clustered to obtain cluster members and midpoints. t-SNE perplexity values and number of clusters $K$ were grid-searched to optimal silhouette scores (i.e. optimal separation of clusters).

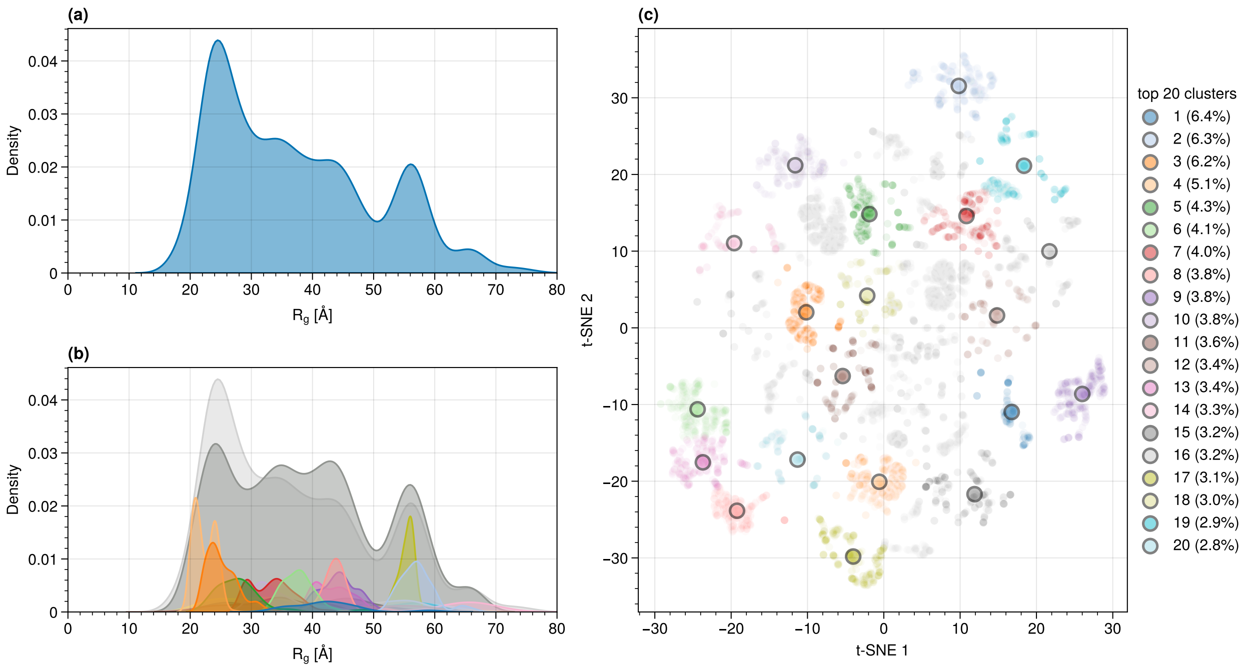


Figure S1: t-SNE clustering structures from the AS trajectory.

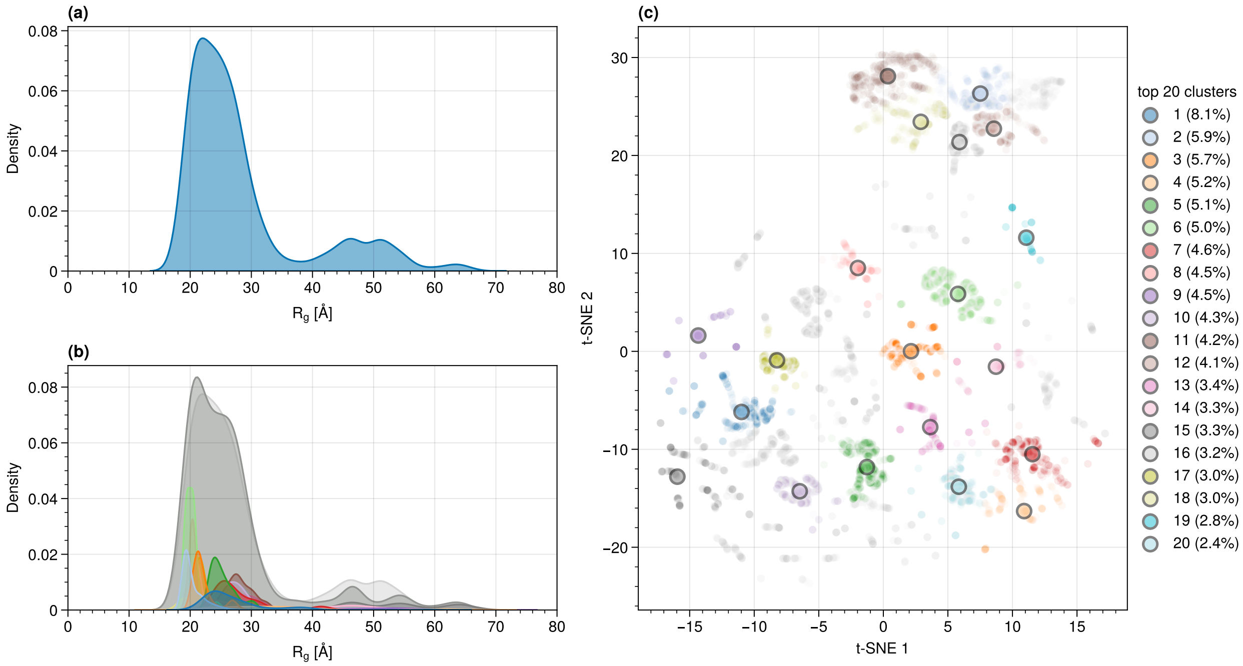


Figure S2: t-SNE clustering structures from the pAS trajectory.

Table S7: Percentage of occurrence of the ten most representative structures of unphosphorylated AS and pAS in the DES-Amber simulation.

| Cluster | AS [%] | pAS [%] |
|---|---|---|
| I | 6.4 | 8.1 |
| II | 6.3 | 5.9 |
| III | 6.2 | 5.7 |
| IV | 5.1 | 5.2 |
| V | 4.3 | 5.1 |
| VI | 4.1 | 5.0 |
| VII | 4.0 | 4.6 |
| VIII | 3.8 | 4.5 |
| IX | 3.8 | 4.5 |
| X | 3.8 | 4.3 |
| | 47.7 | 52.9 |

### 3 Convergence tests

This section investigates the convergence of the simulations of the a99SB-*disp* simulations. The running time averaged values of chemical shifts and secondary structure percentages for AS and pAS are reported in Figure S3(a-b) and (c-d), respectively.

### 4 Calculated Properties and Comparison with Experimental Data

Properties were calculated from the trajectories after the determined convergence of 12 ns. This section covers both the DES-Amber and the a99SB-*disp* force field-based simulations.

**NMR properties.** For each residue in the protein, the shifts chemical of $C_\alpha$, $C_\beta$, N (backbone) and H (backbone) atoms were calculated using ShiftX2.[2] Comparison was made with the experimental values in ref. 3 (Figure S5 and S6, and Table S8).

The prediction of the $^{13}$C-NMR chemical shifts of $C_\alpha$, $C_\beta$ is excellent, even better than that reported in ref. 4 (from a correlation of 0.86 and 0.94 when only using the first 12 residues) for the all residues. The $^{14}$N-NMR also shows a correlation, 0.931 or 0.951, with the experiment. The prediction of $^{1}$H-NMR atoms is less accurate but still acceptable (correlation of 0.295 for a99SB-*disp* and 0.487 for DES-Amber), as seen in.[4]

**CD spectra.** The calculated CD spectrum of AS (Figure S7) is in fair accord as that experimentally measured.[5] The experimental minimum is at a 10 nm lower values than the theoretical value (198 nm and 208 nm, respectively), similarly to what was found in ref. 4.

The minima of the calculated spectra range up to $-15 \cdot 10^{-3}$ deg cm$^2$/dmol for specific conformations, similarly to what found in Rossetti et al., as to be expected given the improvement of the forcefields used,[4] and they average to $-7 \cdot 10^{-3}$ deg cm$^2$/dmol for a99SB-*disp* and $-8 \cdot 10^{-3}$ deg cm$^2$/dmol for DES-Amber, which is in much better agreement with the

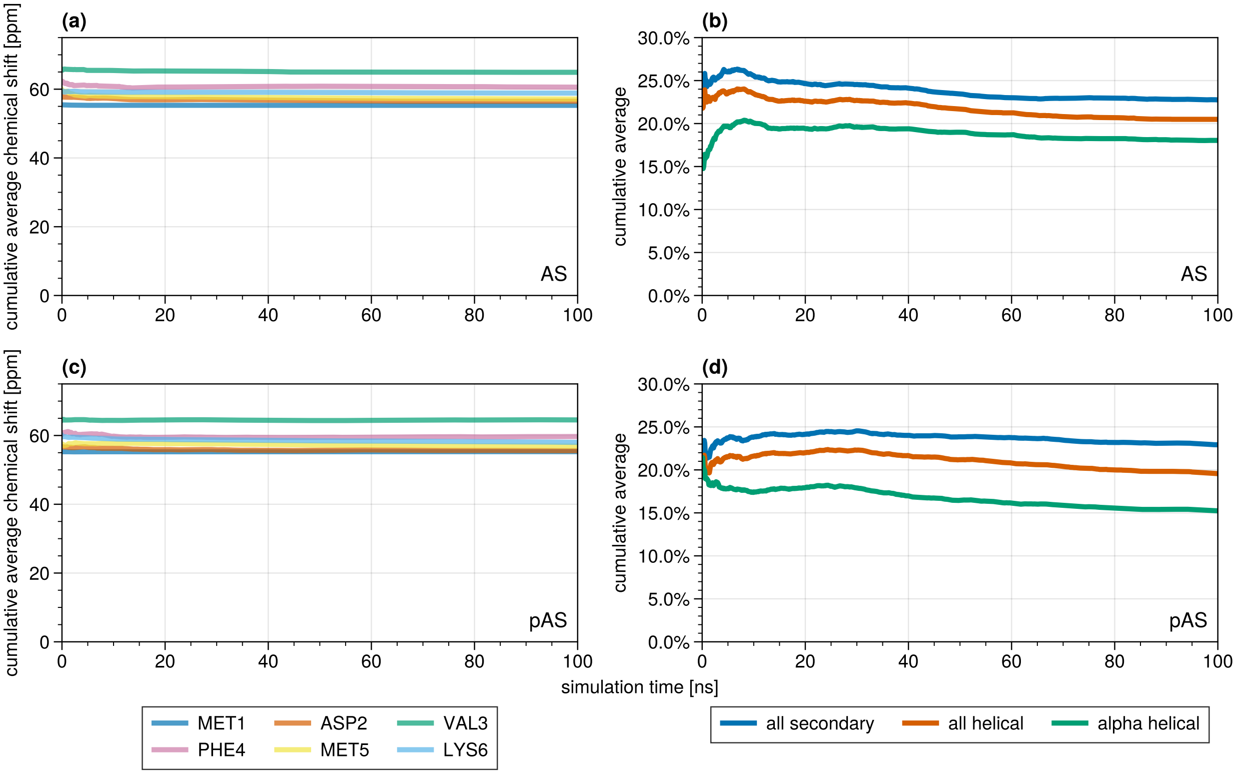


Figure S3: DES-Amber force field-based simulations: (a,c) running time averaged chemical shifts of the $C_\alpha$ atoms of the first six residues; (b,d) running time averaged occurrence of secondary structures (red) and helices (blue) as percent of all residues.

Table S8: Correlation between calculated and measured chemical shifts of *wild-type* AS using both force fields.

| Force field | $^{13}C_\alpha$ | $^{13}C_\beta$ | $^{14}N$ | $^{1}H$ |
|---|---|---|---|---|
| DES-Amber | 0.987 | 0.999 | 0.954 | 0.487 |
| a99SB-*disp* | 0.988 | 0.999 | 0.931 | 0.295 |

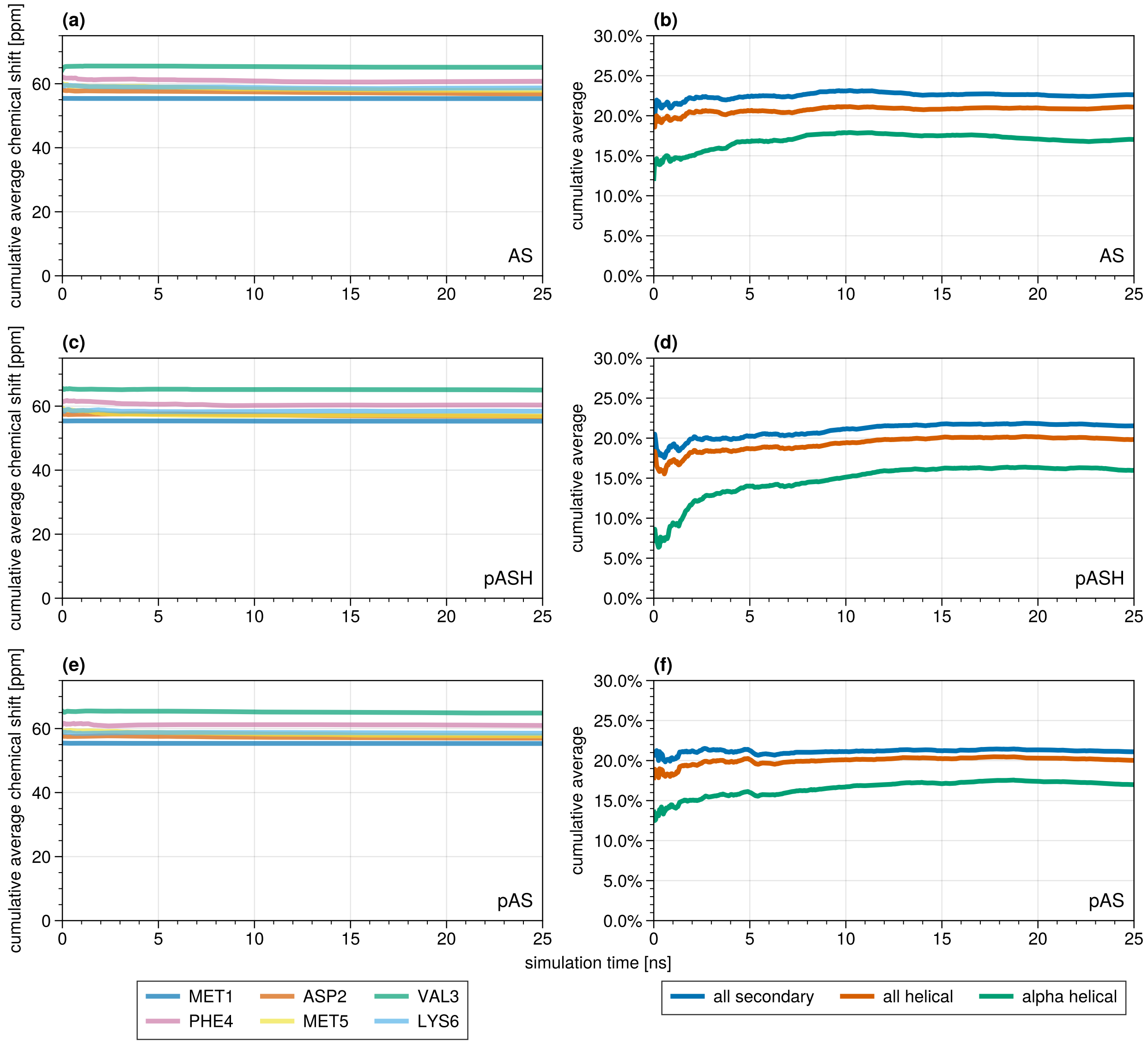


Figure S4: a99SB-*disp* force field-based simulations: (a,c,e) running time averaged chemical shifts of the $C_\alpha$ atoms of the first six residues; (b,d,f) running time averaged occurrence of secondary structures (red) and helices (blue) as percent of all residues.

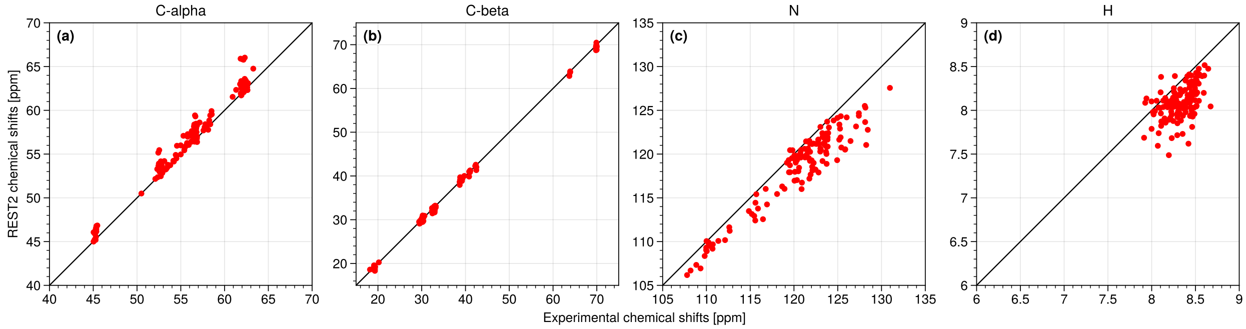


Figure S5: Calculated chemical shifts of $C_\alpha$, $C_\beta$, N and H in AS from the DES-Amber simulation plotted against the experimental data from Roche et al.[3]

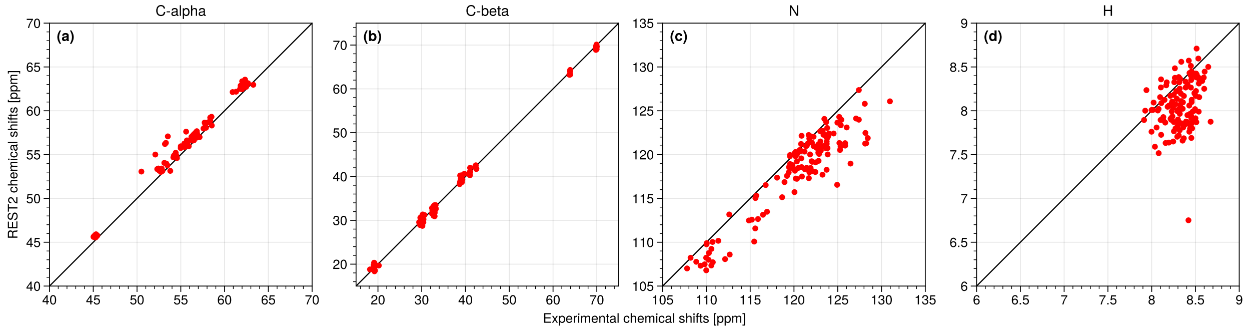


Figure S6: Calculated chemical shifts of $C_\alpha$, $C_\beta$, N and H in AS from the a99SB-*disp* simulation plotted against the experimental data from Roche et al.[3]

experimental results compared to Rossetti et al. The minima in the experimental data of Maltsev et al. are at $-19 \cdot 10^{-3}$ deg cm$^2$/dmol.

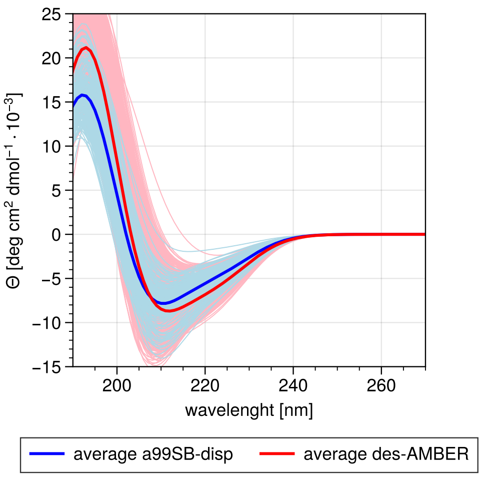


Figure S7: Circular Dichroism spectra of pAS simulations from 12 ns to 25 ns simulation time calculated with DichroCalc;[6] average CD spectrum (red) and CD spectra of all clusters (lightred).

**N- to C-terminus distance.** The distance between the protein termini show no clear trend either in the plot (Figure S8) nor the averages (Table S9).

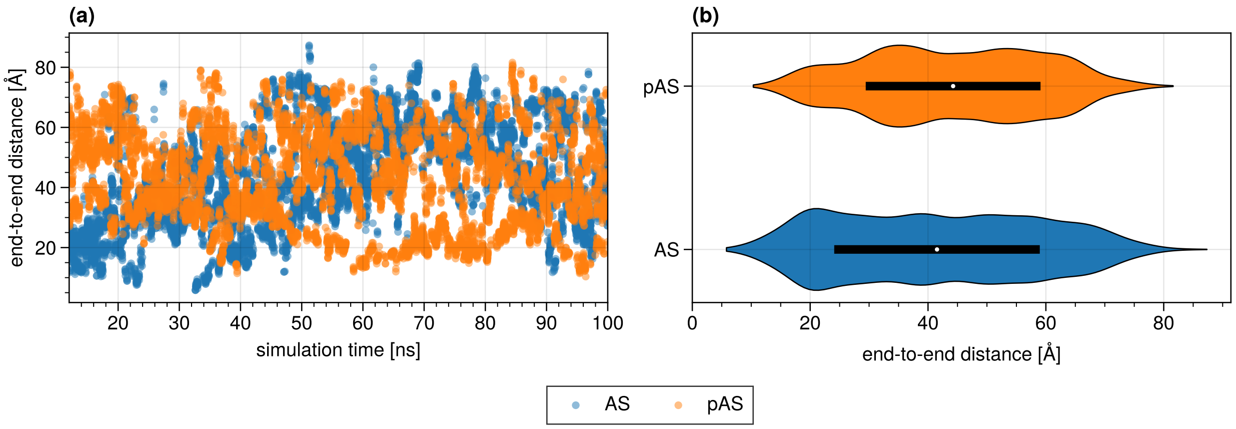


Figure S8: Distance from the N- to C-terminus for both AS and pAS against simulation time in the DES-Amber simulations.

### 5 Amber a99SB-*disp* force field results

Here we present additional data for three secondary simulations using the Amber a99SB-*disp* force field.

**Obtaining structurally similar clusters.**

Table S9: N-terminus – C-terminus distances of AS and pAS averaged over the 12 ns to 100 ns simulation period in the DES-Amber simulations.

| Protein | Distance $d$ [Å] | $\sigma_d$ [Å] |
|---|---|---|
| AS | 44.31 | 16.55 |
| pAS | 43.28 | 15.12 |

Table S10: Number of atoms in the simulation box for the three systems simulated here.

| | Protein | Water | Sodium | Chlorine |
|---|---|---|---|---|
| AS | 2,020 | 68,370 | 75 | 65 |
| pASH | 2,024 | 79,431 | 86 | 75 |
| pAS | 2,023 | 68,361 | 77 | 65 |

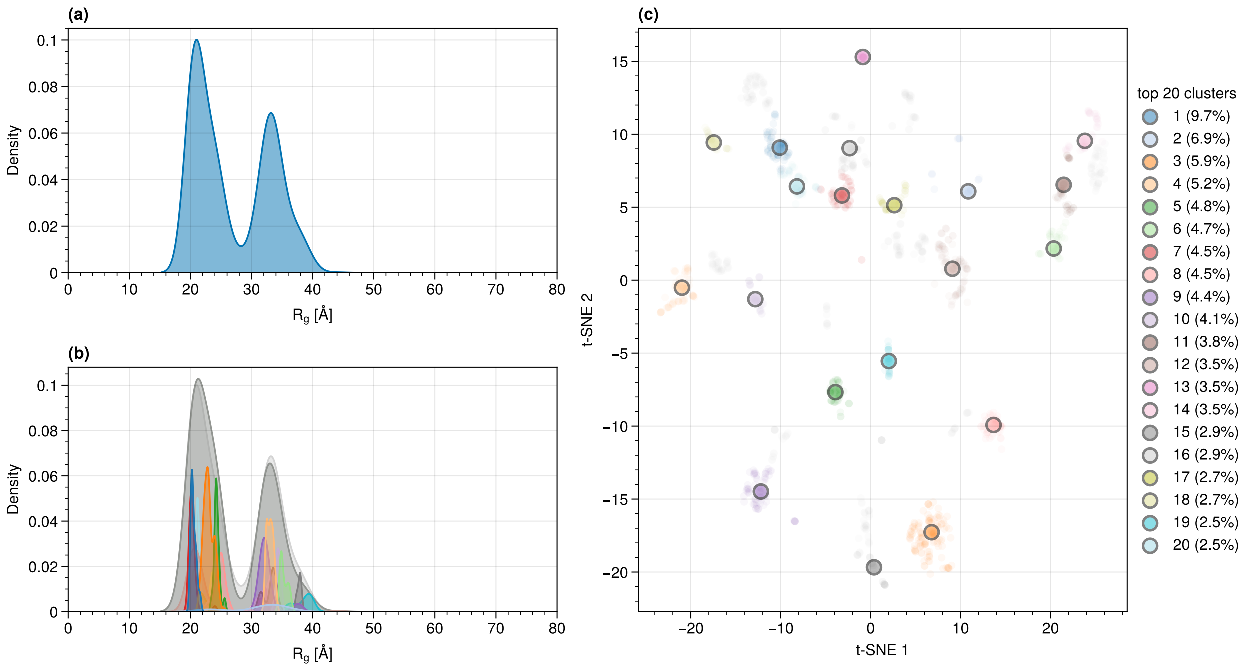


Figure S9: t-SNE clustering structures from the AS trajectory.

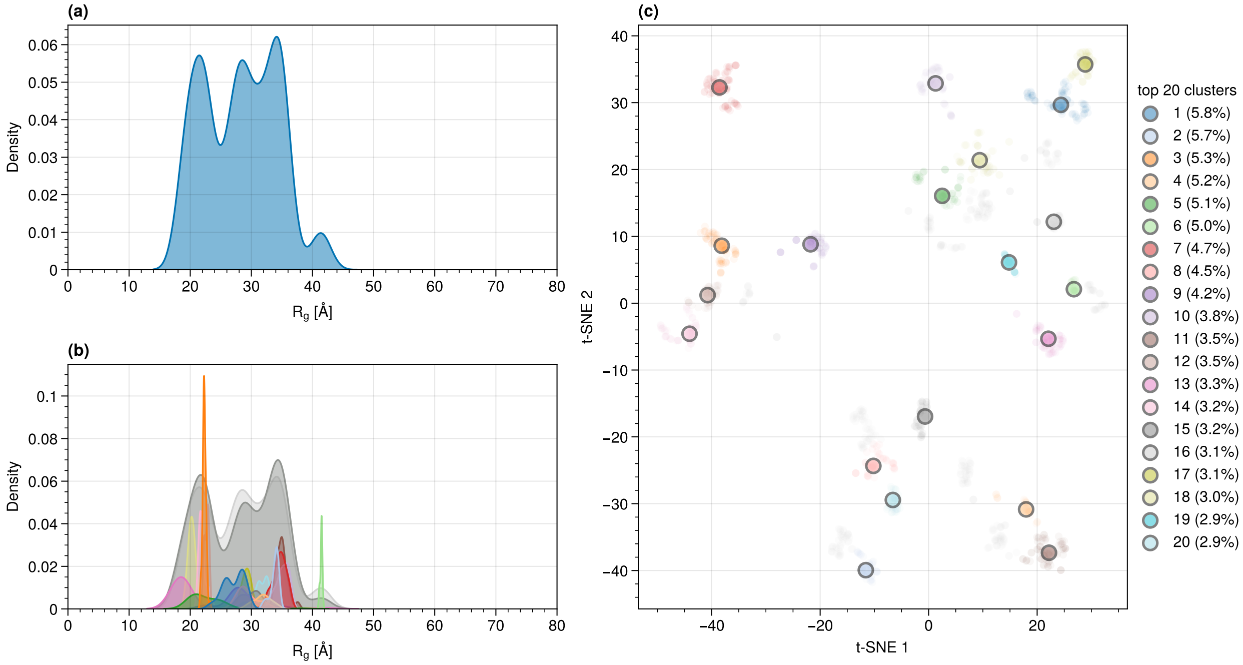


Figure S10: t-SNE clustering structures from the pASH trajectory.

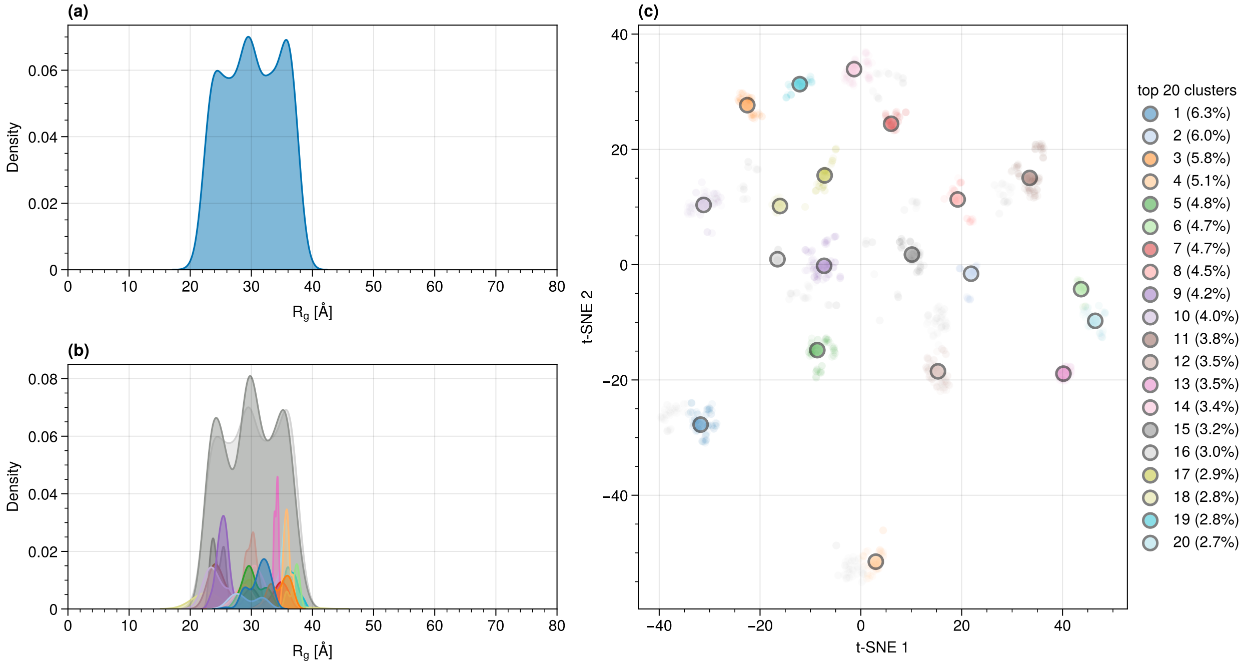


Figure S11: t-SNE clustering structures from the pAS trajectory.

**Size and radius of AS upon phosphorylation.**

Table S11: Calculated properties of AS, pASH and pAS. (i) Hydrodynamic radii $R_H$ and radii of gyration ($R_g$) of the entire proteins and of the hydrophobic regions (HR). The experimental value of $R_H$ is 28.7 Å.[5] (ii) Average number of hydrogen bonds. (iii) Average number of salt bridges. Standard deviations are indicated in parenthesis.

| Protein | $R_H$ [Å] | $R_g$ [Å] | $R_g$(HR) [Å] | $R_H$(HR) [Å] | $N_{SB}$ | $N_{HB}$ |
|---|---|---|---|---|---|---|
| AS | 30.8 (± 3.5) | 27.6 (± 5.7) | 16.7 (± 5.7) | 22.8 (± 5.2) | 2.86 (± 1.96) | 21.51 (± 4.19) |
| pASH | 30.9 (± 3.0) | 27.8 (± 5.5) | 17.4 (± 7.7) | 23.3 (± 6.7) | 1.77 (± 1.53) | 21.02 (± 4.61) |
| pAS | 31.8 (± 2.7) | 29.3 (± 4.5) | 18.7 (± 7.6) | 24.4 (± 6.4) | 1.85 (± 1.62) | 22.13 (± 4.18) |
| AS (exp) | 28.7 | | | | | |

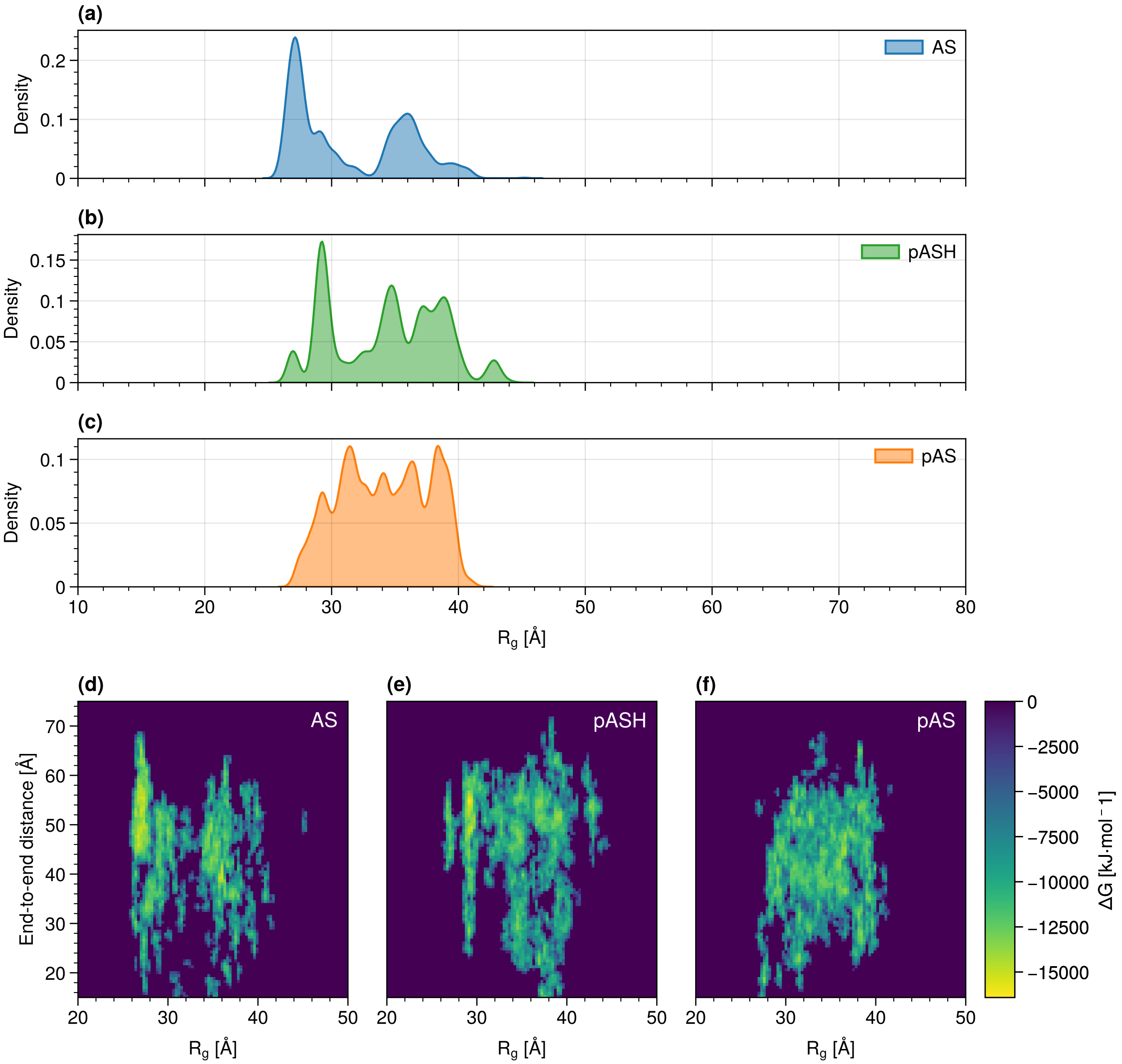


Figure S12: $R_g$ distribution for AS (blue), pASH (green), and pAS (orange) (a-c) and the corresponding (highly approximate) free energy landscapes plotted as a function of the distance between the protein termini (d-f).

**End-to-end distances.** In neither the plot (Figure S13) nor the averages (Table S12) the distance from the N- to C-terminus shows a clear trend.

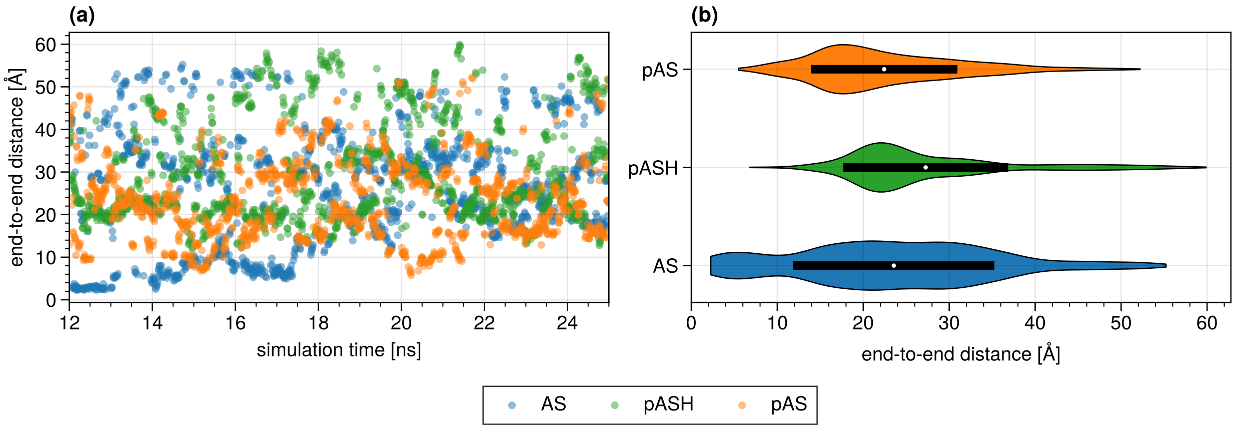


Figure S13: Distance from the N- to C-terminus for all three proteins against simulation time in the a99SB-*disp* simulation.

Table S12: N-terminus – C-terminus distances of AS, pAS, pASH averaged over the 12 ns to 25 ns simulation period in the a99SB-*disp* simulation.

| Protein | Distance $d$ [Å] | $\sigma_d$ [Å] |
|---|---|---|
| AS | 27.8 | 12.9 |
| pASH | 30.8 | 11.9 |
| pAS | 25.9 | 08.6 |

**Solvent accessibility.** The average solvent accessibility of the a99SB-*disp* based simulations (Table S13) closely resemble those found for the simulations based on DES-Amber (Table S13). The averages SASA values are all well within the standard deviation of each other.

Table S13: Average solvent accessible surface areas (SASA) in AS and pAS in the hydrophobic region (HR) and N- and C-terminus using the a99SB-*disp* force field.

| Protein | $\mathrm{SASA_N}$ [Å$^2$] | $\mathrm{SASA_{HR}}$ [Å$^2$] | $\mathrm{SASA_C}$ [Å$^2$] |
|---|---|---|---|
| AS | 87 (± 38) | 83 (± 40) | 107 (± 34) |
| pASH | 90 (± 42) | 84 (± 36) | 112 (± 36) |
| pAS | 86 (± 46) | 85 (± 39) | 107 (± 39) |

**Salt bridges and hydrogen bonds.**

**Ramachandran plot.** The Ramachandran plots (Figure S15) show no discernible differences and exhibit allowed $\phi$ and $\Psi$ values.

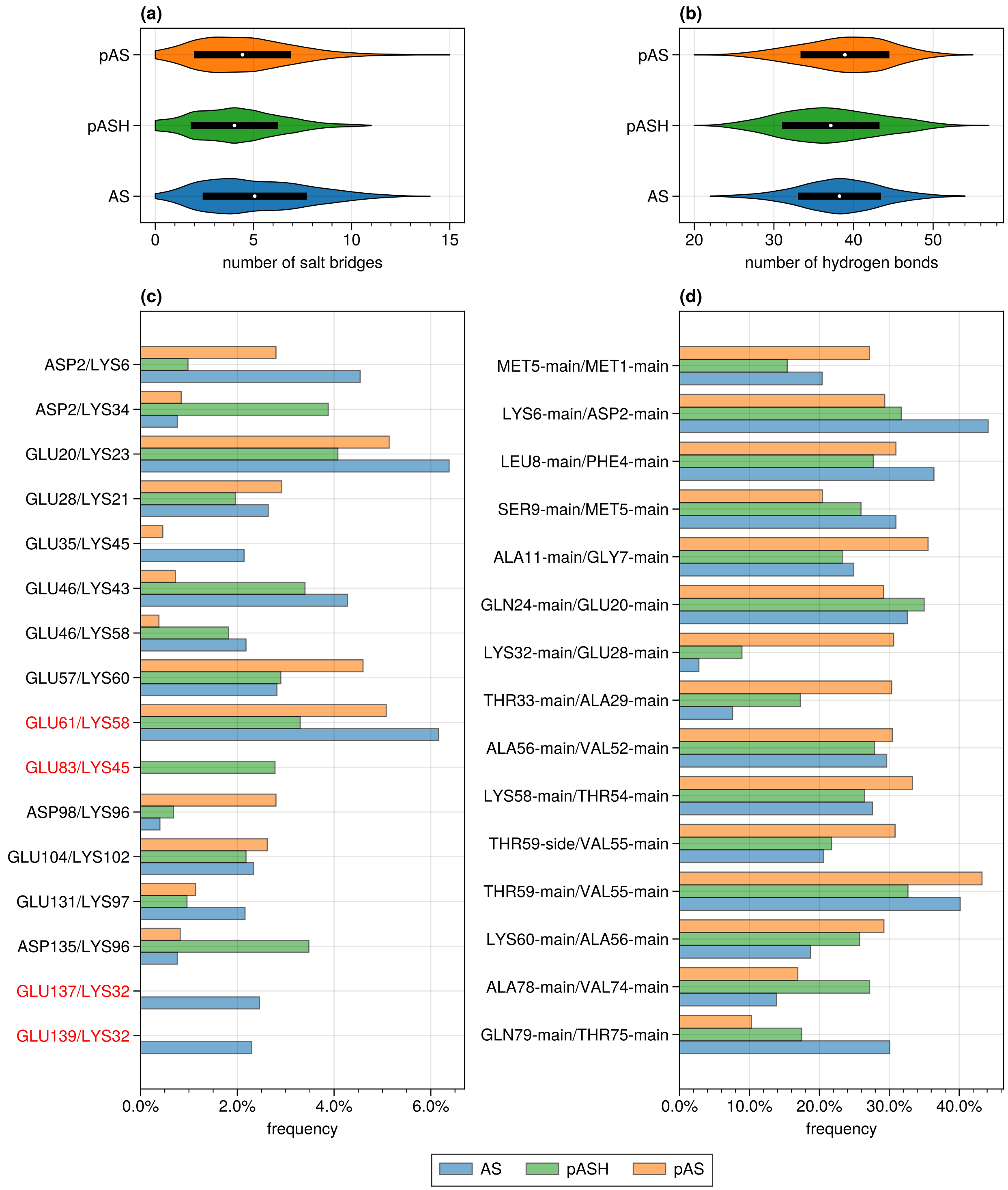


Figure S14: Distribution of the total number of salt bridges (a) and hydrogen bonds (b) in AS (blue), pASH (green) and pAS (orange). Frequency with which intradomain (black labels) and interdomain (red labels) salt bridges (a) and hydrogen bonds (b) are found in AS and pAS. Shown are salt bridges and hydrogen bonds that occur during at least 2 % and 25 % of the converged trajectory, respectively, in either the AS, pASH or pAS simulation.

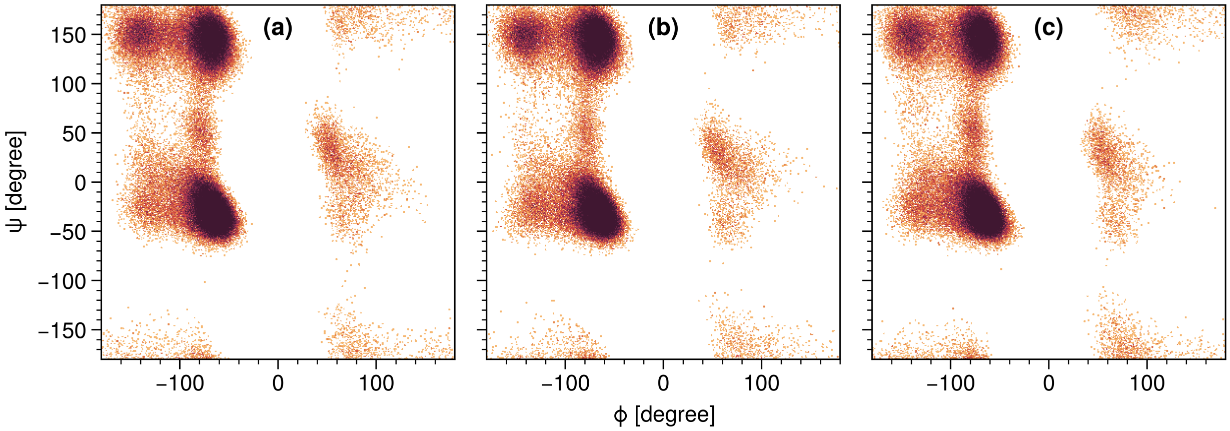


Figure S15: Ramachandran plots for all (a) AS, (b) pASH and (c) pAS from the a99SB-*disp* simulation.

**Radial distribution functions.** This section covers additional RDFs from the a99SB-*disp* simulations.

Some RDFs (Figure S16) did not converge to 1, because the system was not isotropic. The coordination number reads:

$$N(r) = \rho \cdot 4\pi \int_0^r g(r) r^2 \, \mathrm{d}r \tag{1}$$

with $\rho$ being the density of atoms in the simulation box. The integrals are shown up to 0.26 nm (Figure S17) as this covers the peaks visible in Figure S16 which can be attributed to hydrogen bonds.

To eliminate the possibility of double counting solvent molecules, the RDF and integral in Figure S18 show that there are on average 12 solvent molecules surrounding the phosphoryl group in pAS and a slightly reduced 9 around pASH. This is more than the expected maximum of 3 solvent molecules that can be bound via hydrogen bonding per oxygen atom. This was confirmed through visual inspection along the converged part of the trajectories. This over-hydration is attributable to either the high charge density at the site, or an insufficient description of the hydration of phosphoryl moieties by the TIP4P-D water model[7] employed in the current study.

Overall, the a99SB-*disp* force field based simulations show markedly more artefacts in the phosphate hydration than those based on the DES-Amber force field.

### 6 AS with protonated phosphate (pASH)

All results in this section used the a99SB-*disp* force field simulations: following exactly the same protocol as that of pAS.

Both the radius of gyration and hydrodynamic radius (Table S11), the SASA (Table S13) and RDFs (Figure S16) increase on passing from AS to pASH to pAS. The number

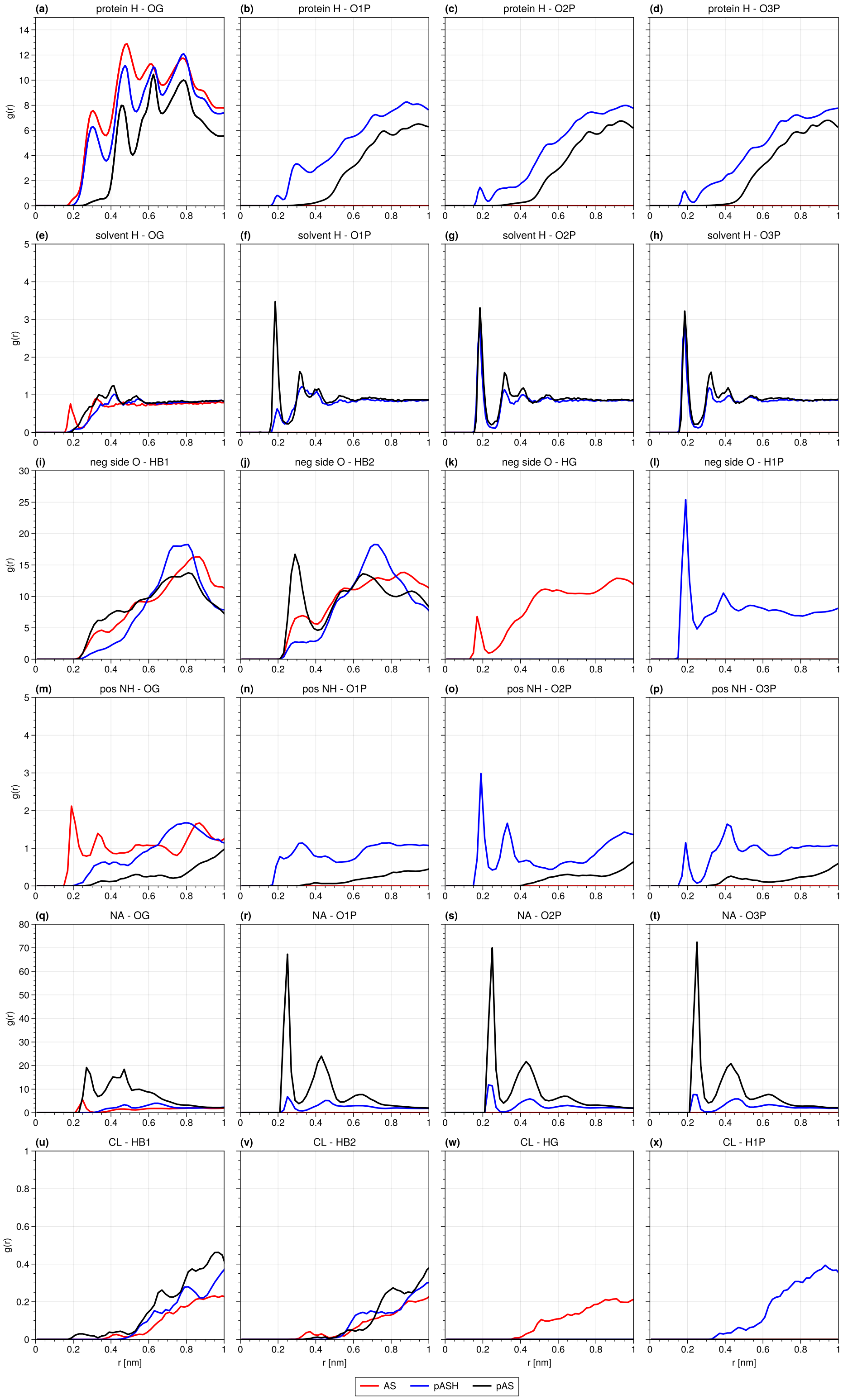


Figure S16: a99SB-*disp* simulation: RDF $g(r)$ of the oxygen atoms of residue 129 side chain and hydrogen atoms bonded to polar atoms of all other residues (a-d), water hydrogens (e-h), hydrogens atoms bonded to positively charged nitrogen (m-p), and sodium ions (q-t) and RDF of hydrogen atoms of residue 129 side chain and negatively charged oxygens of all other residues (i-l) and chlorine ions (u-x). (a), (e), (m), (q) compare the RDF for pAS (black) and AS (red) for the oxygen atom in the sidechain. (b-d), (f-h), (n-p), (r-t) illustrate the rdf for the terminal phosphate oxygens. (i-j), (u-v) compare the rdf of pAS (black) and AS (red) for the sidechain hydrogens bonded to carbon. (k), (w) show the rdf of the hydroxy group. (l), (x) show the rdf of the monoprotonated phosphoryl group.

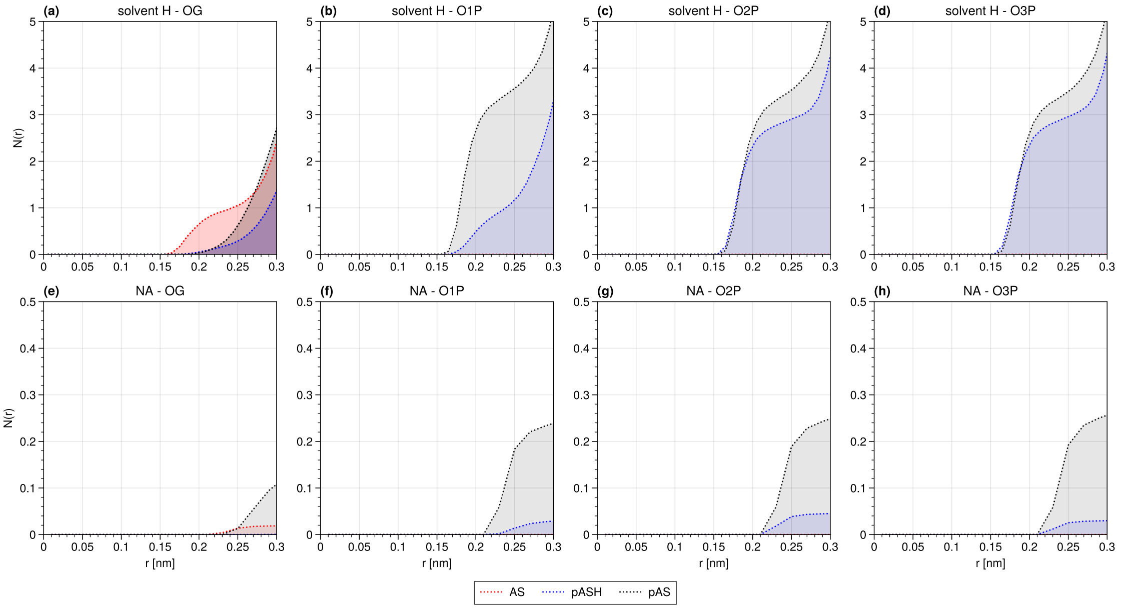


Figure S17: Spherical integrals of the RDFs presented in Figure S16. (a-d) show the integrals of (e-h) in Figure S16 and (e-h) who the integrals of (q-t) in Figure S16. The other integrals are not shown as they are almost zero.

of hydrogen bonds and salt bridges of pASH (Table S11) are lower than those of pAS. The contact maps of pAS and pASH (Figure S19) are similar. The N-terminus/C-terminus distance (Table S9) and the Ramachandran plot (Figure S15) are similar to those of AS and pAS.

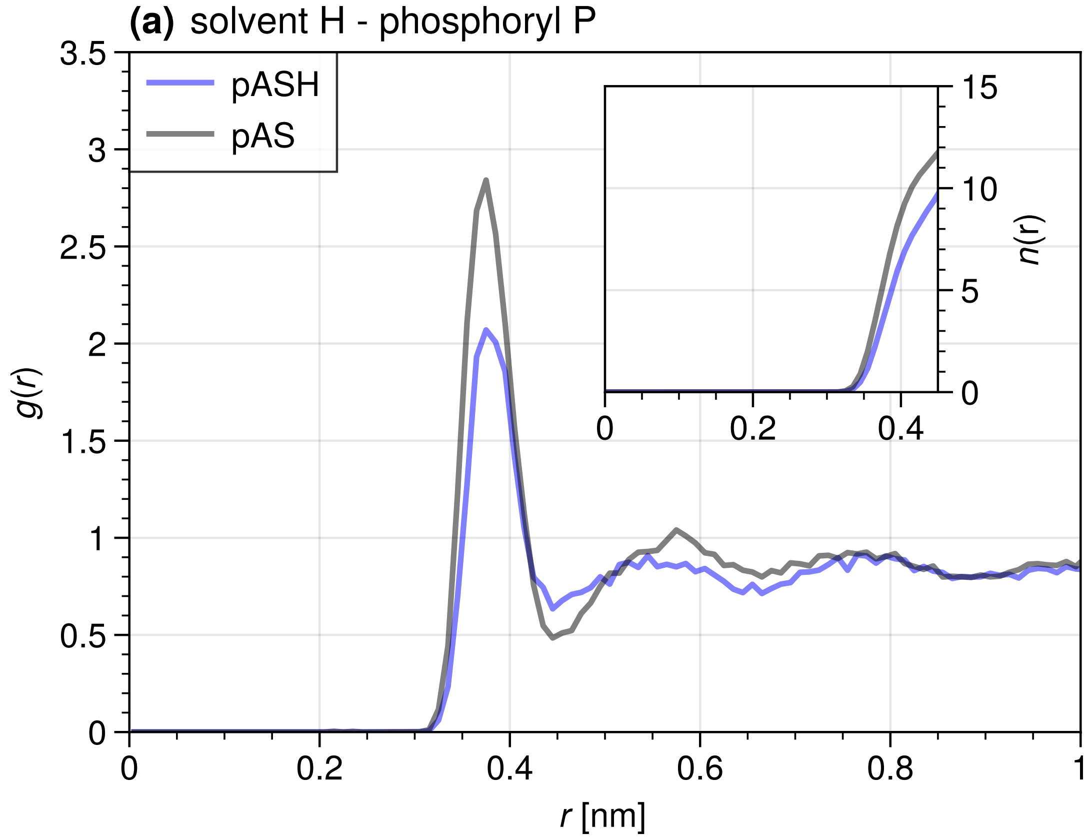


Figure S18: a99SB-*disp* simulation: RDF $g(r)$ of the phosphoryl phosphor atom and solvent hydrogen atoms. The inset shows the cumulative integral $n(r)$ of $g(r)$ up to 4.5 nm (the end of the first solvation shell/peak around the three phosphoryl terminal oxygens).

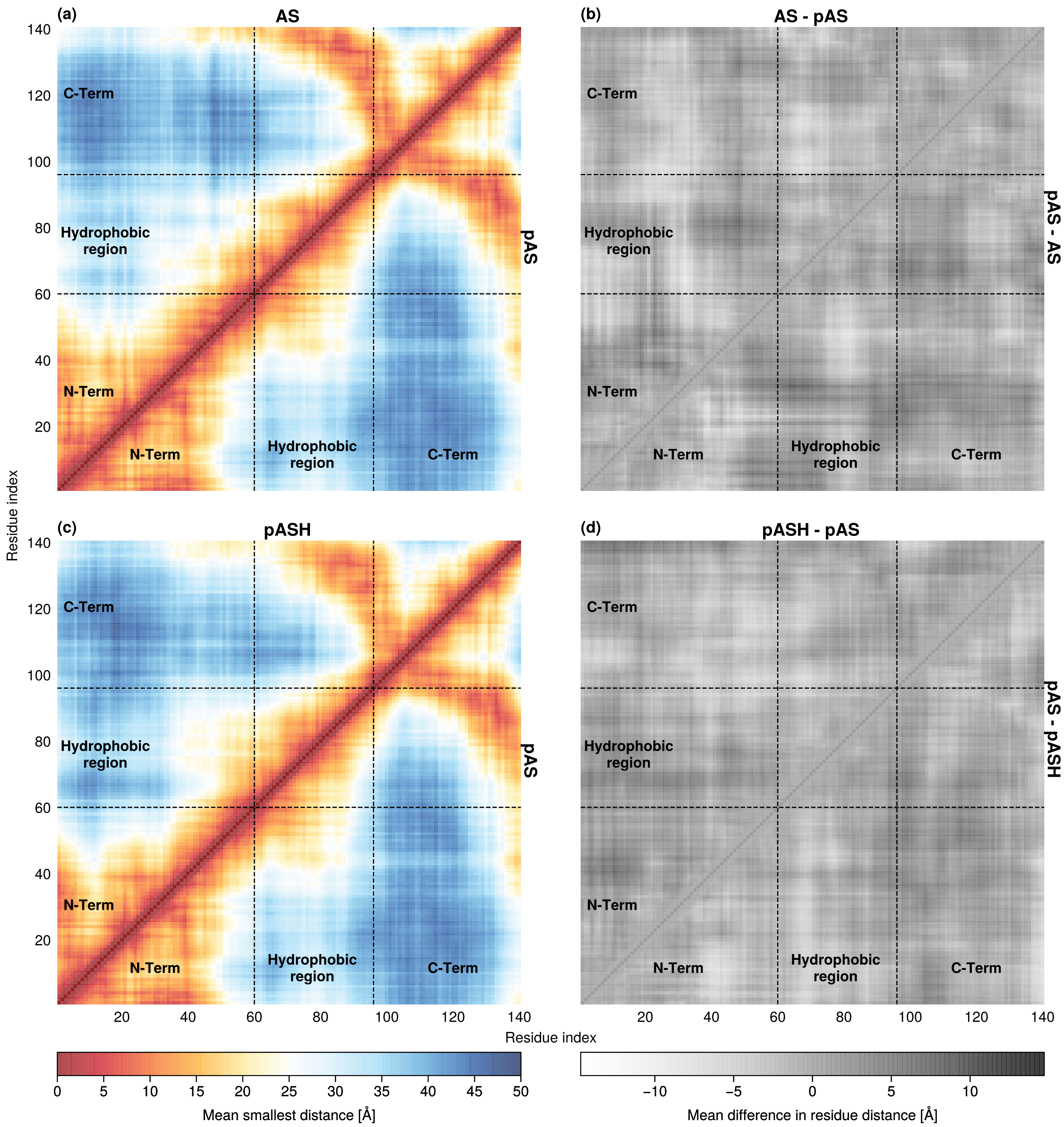


Figure S19: Contact maps of AS (a, triangle above) and pASH (c, triangle above) contrasted with pAS (a,c, triangle below). Mean residue distance differences between AS (b, triangle above) and pASH (d, triangle above) with pAS (b, d, triangle below).

The trajectories of pASH and pAS simulations are broadly similar (Figures S14 to S19). The only exception is in the location of interdomain hydrogen bonds (Figure S14(c)). These differing locations are likely an artefact of insufficient sampling, as the number of hydrogen bonds does not increase with protonation. The phoshporyl group is only slightly less solvated when singly protonated (pASH), compared to its non-protonated form (pAS) (Figure S17). The phosphoryl group at this location is equally highly solvated at this location compared to the unphoshorylated case (AS) and independent of protonation. Indeed, S129 is not involved in salt bridges or hydrogen bonding to any degree in the performed simulations (Figure S14). The contact maps of pASH and pAS (Figure S19) show no significant differences. We conclude that if pASH proteins are present at physiological pH, they contribute to the conformational properties of the protein similarly to those of pAS.

### 7 $\beta$-hairpin-like Structures in the Hydrophobic Region

The converged part of the trajectories were assigned secondary structure designations using the DSSP code.[8,9] The number of loops and irregular elements varies greatly with choice of force field (Figure S20(a,b)). The DES-Amber force field-based simulation exhibit a larger difference in $\alpha$-helix, irregular and loop element, hydrogen bonded turn and bend content than the simulations based on a99SB-*disp* force field. The changes upon phosphorylation in the monomer are more pronounced in the hydrophobic region, regardless of the force field.

We follow ref. 10 to identify possible $\beta$-hairpin-like conformations in the hydrophobic region. The RMSD of a sliding window of 12 residues over the hydrophobic region are calculated with respect to an *ideal $\beta$-hairpin* structure. The $\beta$-hairpin is defined as residues 36-47 of the antimicrobial peptide Arenicin-2 dimer (PDBid: `2L8X`). For each window, the converged part of the simulation trajectories were aligned using the MDAnalysis code.[11–14] We find that the elevated content of $\beta$-hairpin-like structures is highly dependent on the RMSD measure chosen, and depends strongly on the force field chosen (Figure S20(c,d)). Using the same RMSD $\leq 2.5$ Å measure as in ref. 10, the simulations based on DES-Amber display elevated content around residues 61-73. Except for pASH, the simulations based on a99SB-*disp* display notably less content of these structures overall. The pASH simulation also displays higher content of $\beta$-hairpin-like structures around residues 72-84. The content of these structures reduces to around only 2 % when using a slightly tighter criterion of RMSD $\leq 2.0$ Å vs the ideal $\beta$-hairpin.

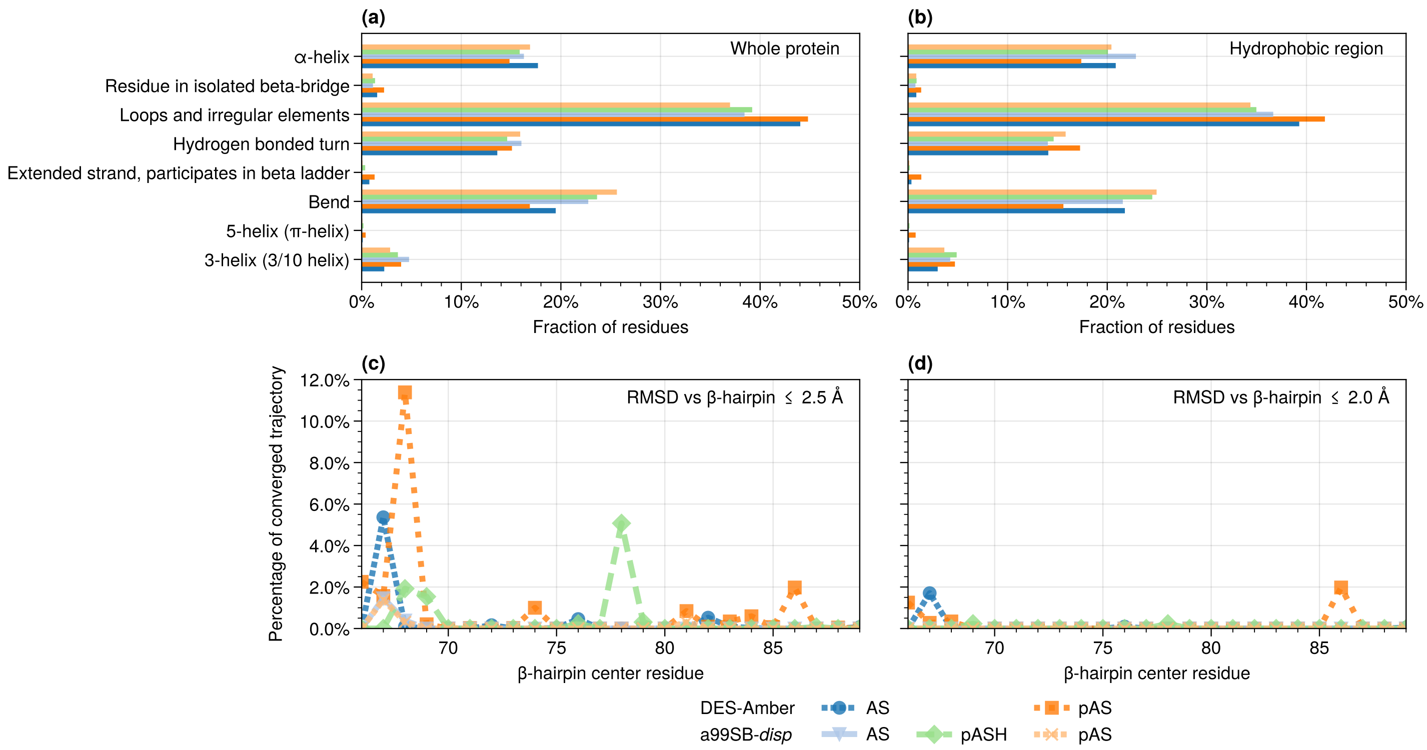


Figure S20: Secondary structure content as found by the DSSP code[8,9] for the whole protein (a) and only the hydrophobic region (b). Percentage of analyzed frames within an RMSD cut-offs of 2.5 Å (c) and 2.0 Å of PDB structure 2L8X for mid-points of 12 residue sliding windows.
